# Comparison of the *in vitro* activity of novel and established nitrification inhibitors applied in agriculture: challenging the effectiveness of the currently available compounds

**DOI:** 10.1101/2020.04.07.023168

**Authors:** Evangelia S. Papadopoulou, Eleftheria Bachtsevani, Eleni Lampronikou, Eleni Adamou, Afroditi Katsaouni, Cécile Thion, Sotirios Vasileiadis, Urania Menkissoglu-Spiroudi, Graeme W. Nicol, Dimitrios G. Karpouzas

**Affiliations:** University of Thessaly, Department of Biochemistry and Biotechnology, Laboratory of Plant and Environmental Biotechnology, Viopolis, 41500 Larissa, Greece; Laboratoire Ampère, École Centrale de Lyon, University of Lyon, 69134 Ecully, France; Aristotle University of Thessaloniki, School of Agriculture, Forestry and Environment, Faculty of Agriculture, Pesticide Science Laboratory, 54124 Thessaloniki, Greece

**Keywords:** nitrification inhibitors, ethoxyquin, quinone imine, ammonia-oxidizing bacteria, ammonia-oxidizing archaea, nitrite-oxidizing bacteria, *in vitro* assays

## Abstract

Nitrification inhibitors (NIs) applied to soil reduce nitrogen fertilizer losses from agricultural ecosystems. Currently available NIs appear to selectively inhibit ammonia-oxidizing bacteria (AOB), while their impact on other groups of nitrifiers is limited. Ethoxyquin (EQ), a preservative shown to inhibit ammonia-oxidizers (AO) in soil, is rapidly transformed to 2,6-dihydro-2,2,4-trimethyl-6-quinone imine (QI) and 2,4-dimethyl-6-ethoxy-quinoline (EQNL). We compared the inhibitory potential of EQ and its derivatives *in vitro* with other established NIs that have been applied in an agricultural setting (dicyandiamide (DCD), nitrapyrin (NP), 3,4-dimethylpyrazole phosphate (DMPP)) by evaluating their impact on the activity and growth of five soil-derived strains (two AOB (*Nitrosomonas europaea, Nitrosospira multiformis*), two ammonia-oxidizing archaea (AOA) (“*Candidatus* Nitrosocosmicus franklandus”, “*Candidatus* Nitrosotalea sinensis”), and one nitrite-oxidizing bacterium (NOB) (*Nitrobacter* sp.)). NIs degradation was also determined. AOA were more sensitive than AOB or NOB to EQ and its derivatives. Despite its transient character, QI was primarily responsible for AO inhibition by EQ, and the most potent NI against AOA. For AOB, QI was more potent than DCD but less than nitrapyrin and DMPP. AOA and NOB showed higher tolerance to the persistent compounds DCD and DMPP. Our findings benchmark the activity range of known and novel NIs with practical implications for their use, and the development of novel NIs with broad or complementary activity against all AO.

---

Modern agricultural systems depend heavily on large inputs of synthetic N fertilizers to maintain crop productivity and alleviate food crisis for the growing global population (1). However, approximately 70% of the annual global input of 100 Tg N fertilizer is lost from agricultural ecosystems due to nitrification and subsequent denitrification processes leading to groundwater and atmospheric pollution through nitrate leaching and nitrogen oxides (NxO) emissions, respectively (2). To reduce N losses and improve nitrogen use efficiency, nitrification inhibitors (NIs) are routinely incorporated into N-stabilized fertilizers to reduce the activities of nitrifying prokaryotes and increase N retention time in soil (3, 4).

Hundreds of compounds have been identified as potential NIs (5), but only three of them have gained importance for practical use on a global scale: 2-chloro-6-(trichloromethyl) pyridine (nitrapyrin) (NP) (6), dicyandiamide (DCD) (7), and 3,4-dimethylpyrazole phosphate (DMPP) (8). All three are known to act on ammonia monooxygenase (AMO), a key enzyme in the first and rate-limiting step of nitrification (9). In particular, NP is believed to act either as copper chelator or “suicide” inhibitor (10, 11), while the mode of action of DCD and DMPP is not yet fully established.

When NIs were first developed for widespread use, nitrification was considered a two-step process carried out by ammonia- (AOB) and nitrite-oxidizing bacteria (NOB). AOB oxidize ammonia to hydroxylamine (NH_2_OH) via AMO, which is further oxidized through to nitrite (NO_2_^-^). NOB subsequently transform NO_2_^-^ to nitrate (NO_3_^-^) via nitrite oxidoreductase (NXR) (12, 13). However, over the last 15 years, other groups contributing to nitrification have been discovered including ammonia-oxidizing archaea (AOA) (14, 15), and ‘comammox’ *Nitrospira* that are able to perform complete oxidation of ammonia to nitrate within an individual cell (16, 17).

These breakthroughs in the microbiology and biochemistry of nitrification were not accompanied by complementary advances on NI research with respect to their spectrum of activity. Most studies have focused on AOB (18–21), and only recently the activity of NIs on AOA was explored (22, 23), while their activity on other groups of nitrifiers (including NOB and comammox bacteria) is not known. Current knowledge of the activity of NIs has been derived from soil microcosm studies where AOB appear to be functionally dominant (24–29). The only available *in vitro* study assessing the comparative activity of NIs on soil-derived AOB and AOA isolates, revealed selective inhibitory activity of DCD and NP against AOB and AOA, respectively (23).

The variation in sensitivity toward different types of NIs, combined with the contribution of AOA, NOB and comammox bacteria to nitrification in distinct ecological niches (30, 31) suggests a suboptimal efficiency of the currently available NIs, and stresses the need for the discovery of novel NIs with a broader range of activity against all microorganisms contributing to nitrification. The use of *in vitro* inhibition assays with a diverse range of soil-derived strains is therefore a necessary benchmarking step to define the exact spectrum of activity of novel and known NIs.

In previous soil microcosm studies we showed that ethoxyquin (EQ) (1,2-dihydro-6-ethoxy-2,2,4-trimethylquinoline), an antioxidant used as preservative in fruit-packaging plants, and its derivative 2,6-dihydro-2,2,4-trimethyl-6-quinone imine (QI), strongly inhibited the activity of AOB and AOA (32). EQ in soil is rapidly transformed to QI and 2,4-dimethyl-6-ethoxyquinoline (EQNL), with the former being the major metabolite (33). The capacity of EQ to be rapidly transformed in soil to possibly potent NIs has particular interest at the application level, considering that the spectrum and the duration of inhibition are desirable attributes of NIs in practice.

We aimed to determine, at the *in vitro* level, the inhibitory potency of EQ and its derivatives on representative soil strains of AOB and AOA, in comparison with three widely used NIs (NP, DCD, and DMPP), whose full spectrum of activity against different ammonia-oxidizers (AO) has not yet been fully established. We expanded our *in vitro* inhibition assays to NOB, whose response to NIs is largely unknown, to gain insights on the impact of NIs on microbial groups functionally associated with ammonia oxidation. Specifically, we determined the inhibitory effects on the growth and activity of two AOB strains (*Nitrosomonas europaea* ATCC25978*, Nitrosospira multiformis* ATCC 25196), two AOA strains (“*Candidatus* Nitrosocosmicus franklandus” C13 and “*Candidatus* Nitrosotalea sinensis” ND2) and one NOB strain (*Nitrobacter* sp. NHB1) using q-PCR to measure the abundance of *amoA* or *nxrB* functional genes, and the production or consumption of nitrite, for AO or NOB, respectively. To establish possible correlations between NI presence and inhibition, their degradation and transformation (for EQ) were also determined.

## RESULTS

### The impact of NIs on the activity and growth of AOB isolates

The effects of six NIs (EQ, QI, EQNL, DCD, NP and DMPP) on the activity and growth of the two AOB isolates were initially tested over a range of concentration levels. EQ strongly inhibited ammonia oxidation in liquid cultures of *N. europaea* (Fig. 1a) and *N. multiformis* (Fig. 2a) only at the concentration of 460 μM. QI completely inhibited ammonia oxidation by *N. europaea* (Fig. 1b) and *N. multiformis* (Fig. 2b) at concentrations ≥ 270 μM and ≥ 135 μM, respectively. In contrast, EQNL only temporarily inhibited *N. europaea* activity at the highest concentration tested, 500 μM (Fig. 1c), while at the same concentration level *N. multiformis* activity was fully inhibited (Fig. 2c).

**Fig. 1.**
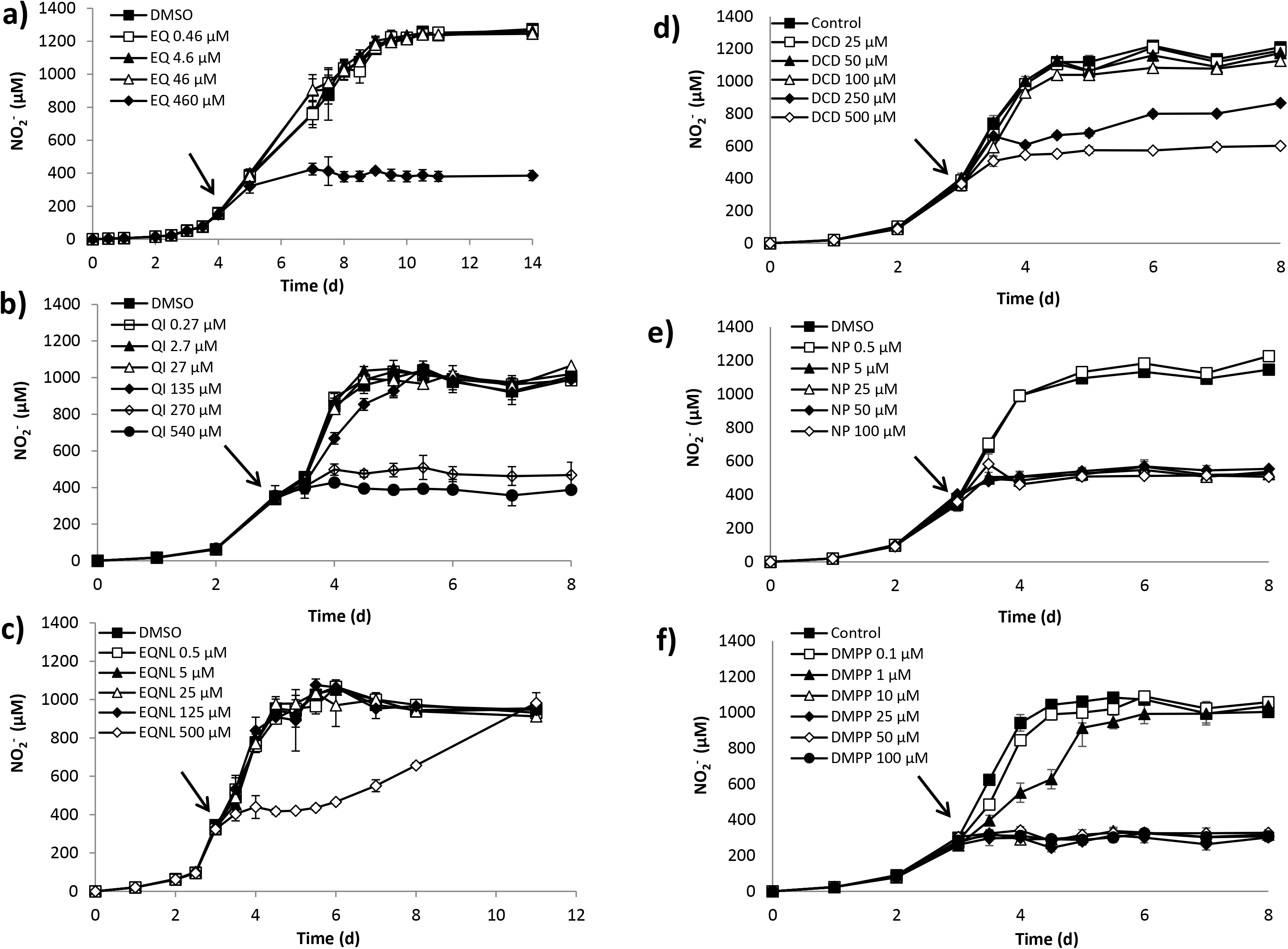
The effect of different concentrations of Ethoxyquin (EQ) (a), Quinone Imine (QI) (b), Ethoxyquinoline (EQNL) (c), Dicyandiamide (DCD) (d), Nitrapyrin (NP) (e), and DMPP (f), on nitrite production by *Nitrosomonas europaea*. Error bars represent the standard error of the mean of triplicate cultures. Arrows indicate the time point when the nitrification inhibitor (NI) was added.

**Fig. 2.**
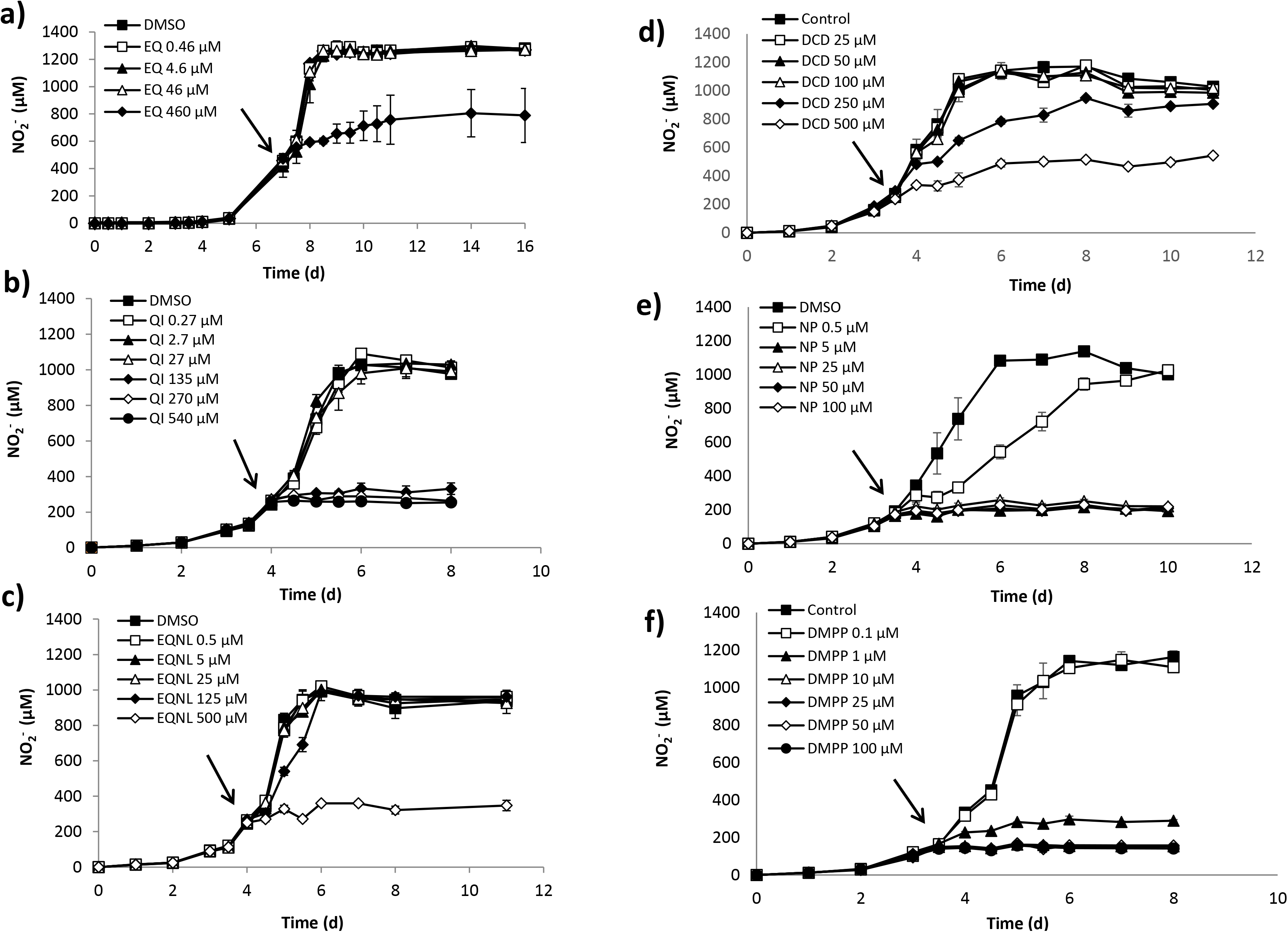
The effect of different concentrations of Ethoxyquin (EQ) (a), Quinone Imine (QI) (b), Ethoxyquinoline (EQNL) (c), Dicyandiamide (DCD) (d), Nitrapyrin (NP) (e), and DMPP (f) on nitrite production by *Nitrosospira multiformis*. Error bars represent the standard error of the mean of triplicate cultures. Arrows indicate the time point when the nitrification inhibitor (NI) was added.

For the established NIs, DCD inhibited the activity of both AOB strains at concentrations of 250 μM and 500 μM (Fig. 1d), with a mere recovery of *N. multiformis* at 250 μM (Fig 2d). NP completely inhibited the activity of both *N. europaea* and *N. multiformis* at concentrations of ≥ 5 μM (Fig. 1e and 2e). Similarly, DMPP reduced nitrite production by both AOB isolates at concentrations ≥1 μM, with a late recovery observed only for *N. europaea* at 1 μM (Fig. 1f and 2f), and complete inhibition observed at concentrations ≥10 μM. The inhibition of AOB growth as determined by measurements of *amoA* gene abundance were in agreement with the NO_2_^-^ production rates (Fig. S1 and S2) with the exception of a low level inhibition (22.9 ± 3.9 %) of *N. multiformis* growth at 0.1 μM DMPP compared to the control, despite no difference in nitrite production rates (Fig. S2f).

Based on the inhibition assays, EC_50_ values for each strain x compound combination were calculated. The two AOB showed equivalent EC_50_ values for the different NIs tested with the sole exception of EQ derivatives. Higher EC_50_ values were observed for *N. europaea* (199.8±16.0 and 543.4±111.5 μM for QI and EQNL, respectively) compared to *N. multiformis* (65.1±6.O and 36O.5±1O5.7 μM) (Table 1). DMPP and NP were the most potent inhibitors of *N. europaea* (EC_50_ = 2.1±0.7 and 2.1±O.4 μM, respectively), followed by EQ, QI, and DCD which did not differ in their ability to inhibit *N. europaea*, with EQNL being the weakest inhibitor (EC_50_ = 181.4±23.3 μM) (Table 1). For *N. multiformis*, DMPP, NP, and QI were equally effective and the most potent inhibitors, with EC_50_ values of O. 6±O.1, O.8±O.3, and 65.1±O.6 μM, respectively, followed by EQ (EC_50_ = 214.8±39.6 μM), DCD (EC_50_ = 248.7±7.4 μM) and EQNL (EC_50_ = 36O.5±1O5.7 μM) (Table 1).

**Table 1.**
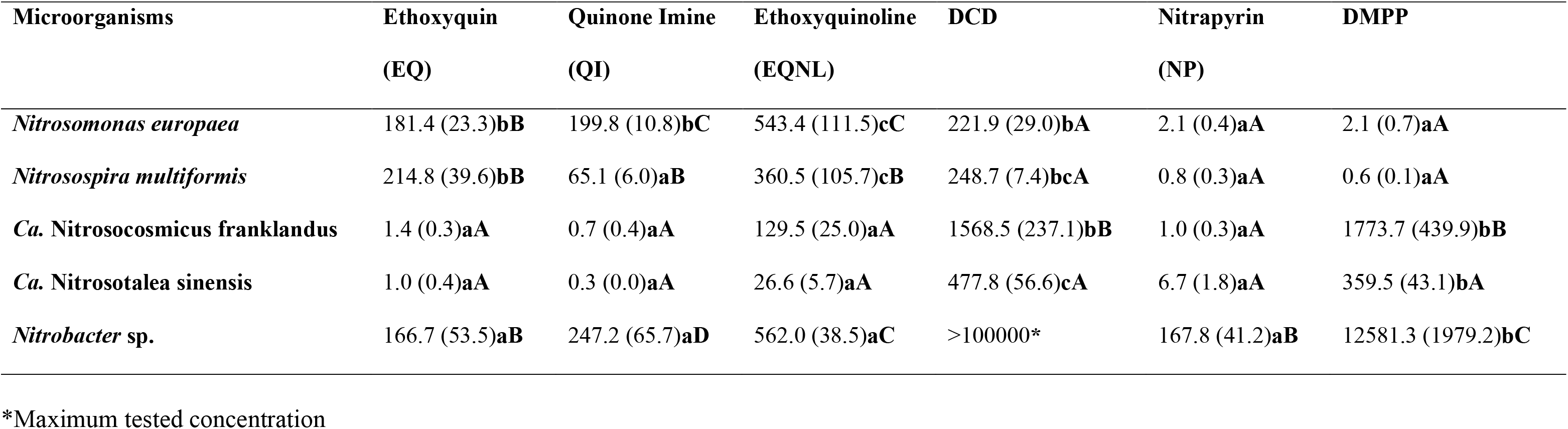
Mean EC_50_ values (μM) of the tested nitrification inhibitors (NIs) on ammonia or nitrite oxidation activity of the nitrifying strains. Standard errors of the mean values are given in brackets. Upper case letters indicate significant differences (p<0.05) between microorganisms for each individual NI, and lower-case letters indicate significant differences (p<0.05) between NIs for each tested microorganism.

### The impact of NIs on the activity and growth of AOA isolates

We further tested the inhibitory effects of NIs on the activity and growth of two soil-derived AOA isolates. The activity of “*Ca*. N. franklandus” (Fig. 3a) and “*Ca*. N. sinensis” (Fig. 4a) was significantly reduced by EQ at concentrations ≥ 4.6 μM and ≥ 0.46 μM, respectively. However, complete recovery of “*Ca*. N. sinensis” activity was observed at 0.46 μM (Fig. 4a). QI significantly reduced the activity of AOA at all concentrations, with a gradual recovery observed for only “*Ca*. N. franklandus” at the lowest concentration level (0.27 μM) (Fig. 3b and 4b). EQNL suppressed ammonia oxidation by “*Ca*. N. franklandus” (Fig. 3c) and “*Ca*. N. sinensis” (Fig. 4c) at concentrations ≥ 125 μM and ≥ 25μM, respectively, although recovery was observed. The impact of EQNL on the growth of “*Ca*. N. franklandus” was not fully consistent with the activity measurements, and no significant differences among the different concentrations were observed at the end of the incubation period (Fig. S3).

**Fig. 3.**
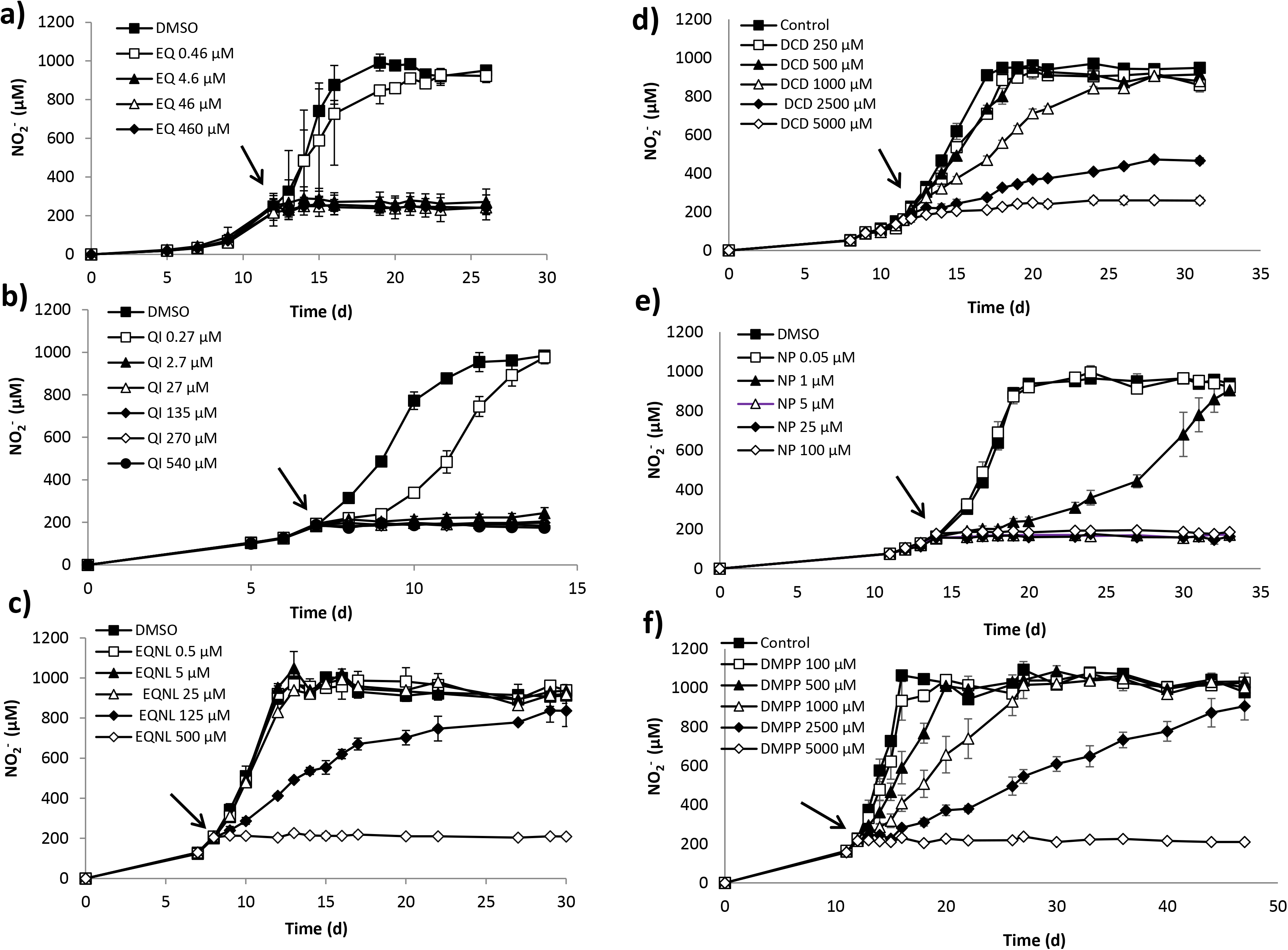
The effect of different concentrations of Ethoxyquin (EQ) (a), Quinone Imine (QI) (b), Ethoxyquinoline (EQNL) (c), Dicyandiamide (DCD) (d), Nitrapyrin (NP) (e), and DMPP (f) on nitrite production by “*Candidatus* Nitrosocosmicus franklandus”. Error bars represent the standard error of the mean from triplicate cultures. Arrows indicate the time point when the nitrification inhibitor (NI) was added.

**Fig. 4.**
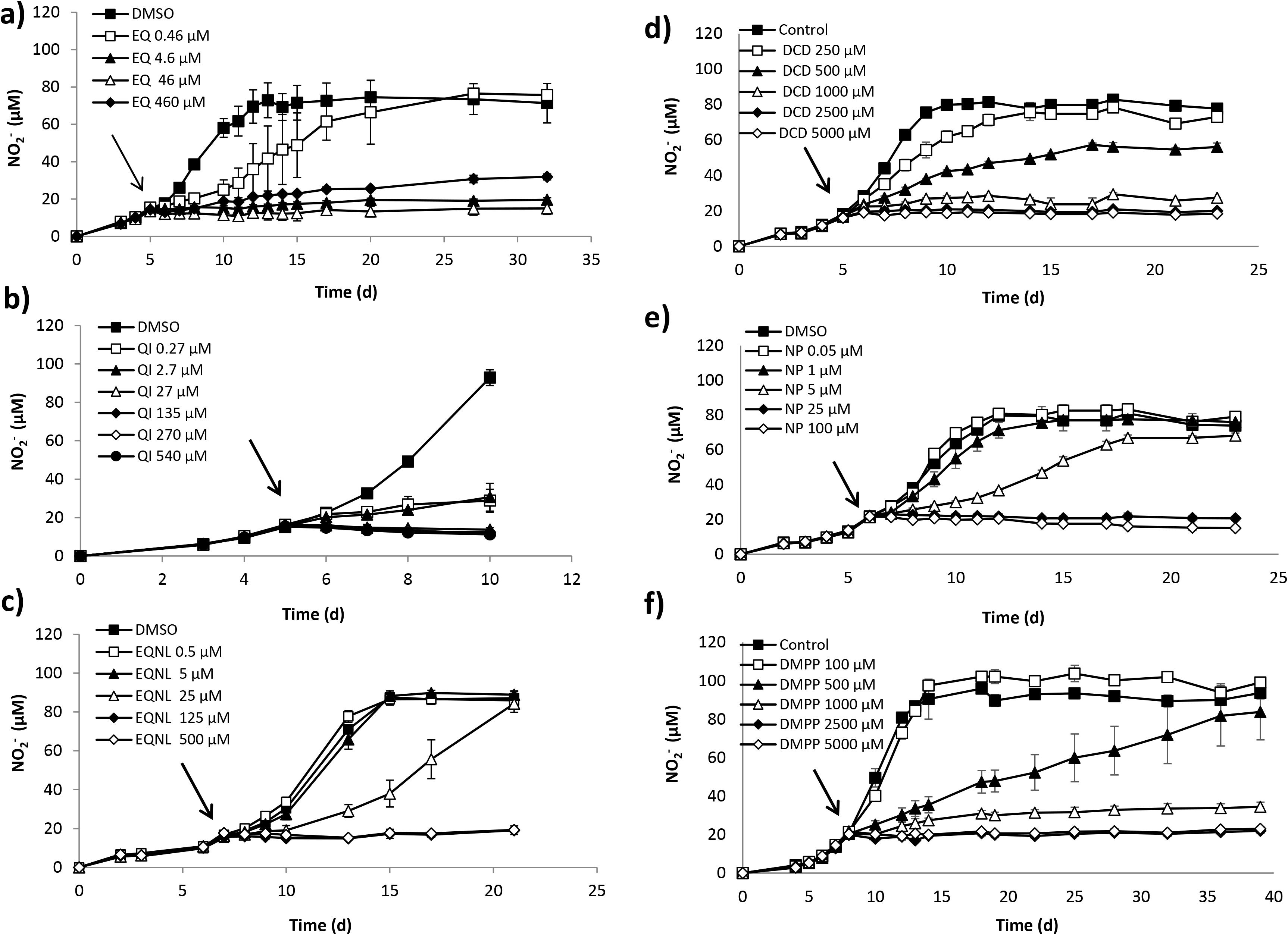
The effect of different concentrations of Ethoxyquin (EQ) (a), Quinone Imine (QI) (b), Ethoxyquinoline (EQNL) (c), Dicyandiamide (DCD) (d), Nitrapyrin (NP) (e), and DMPP (f) on nitrite production by “*Candidatus* Nitrosotalea sinensis”. Error bars represent the standard error of the mean of triplicate cultures. Arrows indicate the time point when the nitrification inhibitor (NI) was added.

Regarding the other NIs, DCD halted the activity of “*Ca*. N. franklandus” (Fig. 3d) and “*Ca*. N. sinensis” (Fig. 4d) at concentrations ≥ 1000 μM and ≥ 500 μM respectively, with a complete recovery observed only for “*Ca*. N. franklandus” at 1000 μM. However, based on growth measurements, DCD significantly reduced “*Ca*. N. franklandus” growth even at 500 μM (Fig. S3). NP significantly reduced the activity of “*Ca*. N. franklandus” (Fig. 3e) and “*Ca*. N. sinensis” (Fig. 4e) at concentrations ≥ 1μM and ≥ 5μM, respectively, although recovery was observed at these concentrations at the end of the incubation. Finally, DMPP inhibited the activity of both AOA isolates at concentrations ≥ 500 μM (Fig. 3f and 4f). However, complete recovery was observed for “*Ca*. N. franklandus” at 500, 1000, and 2500 μM (Fig. 3f), and for “*Ca*. N. sinensis” only at 500 μM (Fig. 4f). In certain cases, the impact of DMPP on nitrite production was not concomitant with growth patterns. Hence, DMPP concentrations ≥ 500 μM induced a persistent reduction in *amoA* gene abundance of “*Ca*. N. franklandus” (Fig. S3), while “*Ca*. N. sinensis” showed complete recovery of its growth at all tested concentrations at the end of the incubation period (Fig. S4).

When EC_50_ values were calculated “*Ca*. N. franklandus” differed in its sensitivity to DCD and DMPP compared to “*Ca*. N. sinensis”, with the former being more tolerant to both DCD (EC_50_ of 1568.5±237.1 μM compared to 477.8±56.6 μM for “*Ca*. N. sinensis”), and DMPP (EC_50_ of 1773.7±359.5 μM compared to 359.5.5±43.1 μM for “*Ca*. N. sinensis”) (Table 1). DCD and DMPP were also the weakest AOA inhibitors from those tested (Table 1). In contrast, EQ and its oxidative derivatives as well as NP were equally effective inhibitors of both AOA isolates, with QI scoring the lowest EC_50_ values (0.7±0.4 and 0.3±0.0 μM for “*Ca*. N. franklandus” and “*Ca*. N. sinensis”, respectively).

### The impact of NIs on the activity and growth of *Nitrobacter* sp. NHB1

Nitrite oxidation by *Nitrobacter* sp. NHB1 was completely inhibited by EQ, EQNL, and DMPP only at the highest tested concentrations of 460 μM, 500 μM, and 25000 μM, respectively, and DCD was not inhibitory even at the highest concentration tested (100000 μM) (Fig. 5). In contrast, QI and NP significantly suppressed *Nitrobacter* sp. NHB1 activity at concentrations ≥ 135 μM and ≥1OO μM, respectively, though recovery was observed for QI at 135 μM and 270 μM (Fig. 5). We observed variations in the growth inhibition patterns of the tested NIs (assessed as a lack of increase in *nxrB* gene abundance). EQ induced a significant reduction in the growth of *Nitrobacter* sp. NHB1 at all concentration levels at the end of the incubation period. QI strongly inhibited growth at concentrations ≥135 μM, while EQNL did not affect bacterial growth even at the highest concentration level (500 μM) (Fig. S5). On the contrary, DCD at 100000 μM, and DMPP at concentrations ≥ 25000 μM significantly suppressed *Nitrobacter* sp. NHB1 growth (Fig. S5). When EC_50_ values were calculated EQ, QI, EQNL and NP were equally suppressive towards *Nitrobacter* sp. NHB1, while DMPP and DCD showed no appreciable inhibition (Table 1).

**Fig. 5.**
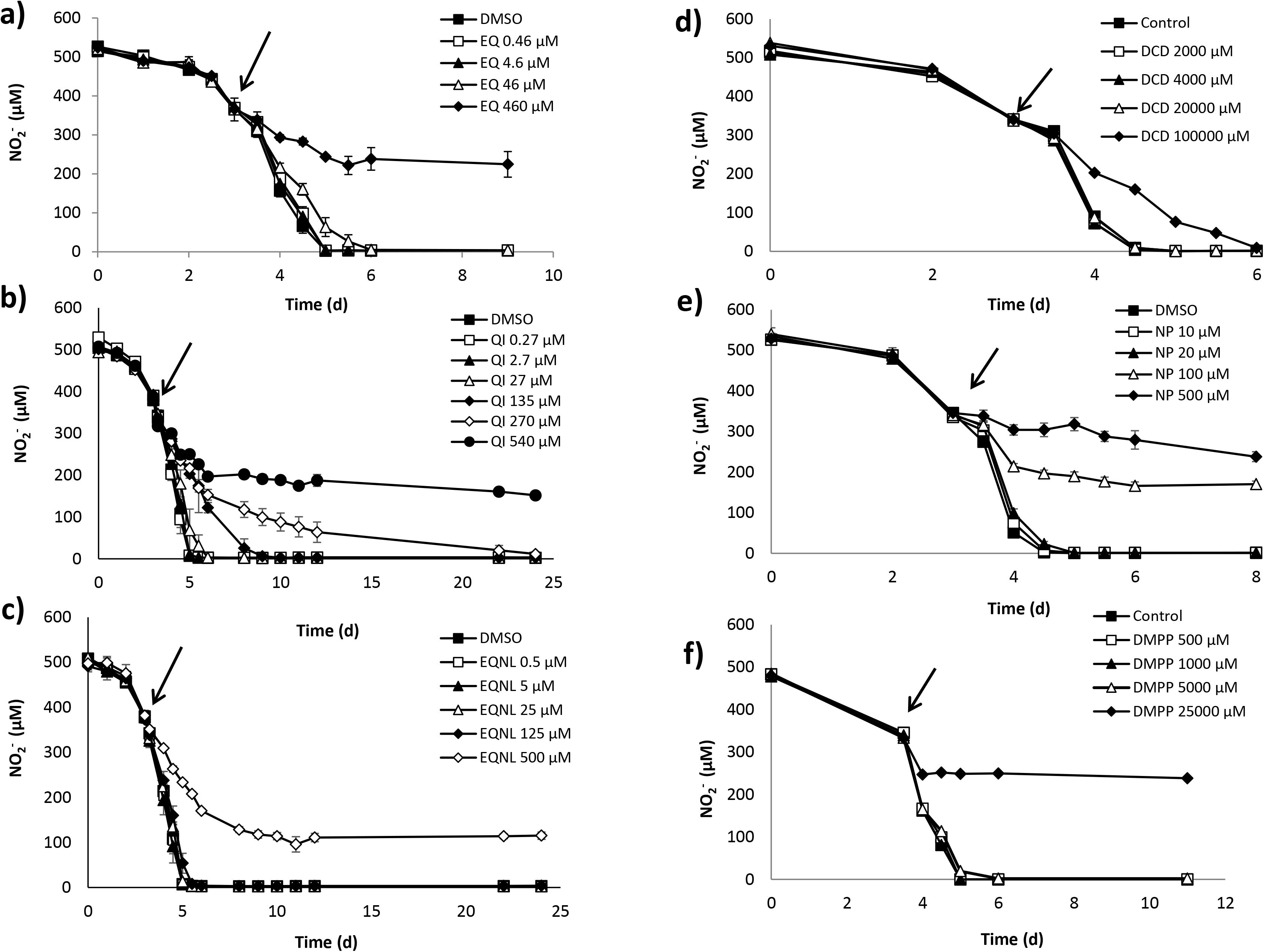
The effect of different concentrations of Ethoxyquin (EQ) (a), Quinone Imine (QI) (b), Ethoxyquinoline (EQNL) (c), Dicyandiamide (DCD) (d), Nitrapyrin (NP) (e), and DMPP (f) on nitrite transformation by *Nitrobacter* sp. Error bars represent the standard error of the mean from triplicate cultures. Arrows indicate the time at which the nitrification inhibitor (NI) was added.

### Degradation of NIs in the *in vitro* assays

We further determined the degradation of all tested NIs during the *in vitro* assays to establish whether there is a relationship between the length of exposure and inhibition. The degradation of NIs in most cases was best described by the single first order (SFO) kinetic model (*x*^2^ ≤15, *r*^2^ ≥O.75), with the exception of NP which in certain treatments followed a biphasic degradation pattern (Table S1). EQ was rapidly transformed to QI and EQNL. QI was detected at concentrations below its EC_50_ values for AOB/NOB but above its EC_50_ values for AOA at the onset of EQ-induced inhibition, (Tables 1 and 2). However, the maximum concentrations of QI formed in these cultures were equivalent or higher than the EC_50s_ for *N. europaea* and *N. multiformis* (AOB), respectively, while still lower than that for *Nitrobacter* sp. NHB1 (NOB). In contrast, EQNL was always formed at much lower concentrations than the estimated EC_50_s for AO and NOB with the exception of its maximum recorded concentration in the liquid culture of “*Ca*. N. sinensis” amended with 460 μM EQ (Table 1 and 2, Fig. S6). The degradation half-life (DT50) for the sum of EQ+QI+EQNL ranged from 2.1 to 60.1 days in the cultures of *Nitrobacter* sp. NHB1 and *N. multiformis* respectively when amended with 460 μM of EQ (Table S1). QI showed limited persistence and a weak dose-dependent degradation pattern with DT50 = 0.05 - 1.52 days at the lowest concentration level (2.7 μM) and 2.23 - 5.65 days at the highest concentration level (540 μM). In contrast, EQNL persisted in the liquid cultures throughout the experiment (extrapolated DT50 values >1000 days) (Table S1).

**Table 2.** Mean concentrations ± standard errors (μM) of Quinone Imine (QI) and Ethoxyquinoline (EQNL) formed in the liquid cultures of the nitrifying isolates amended with Ethoxyquin (EQ) (i) at the onset of inhibition, (ii) at the time when maximum concentration levels were detected. The timepoint (days) at which each measurement was taken is given in brackets.

| Ethoxyquin treatment | | Concentration ($\mu\text{M}$) | | | |
| --- | --- | --- | --- | --- | --- |
|  |  | Inhibition Onset |  | Maximum concentration formed |  |
|  |  | QI | EQNL | QI | EQNL |
| **AOB** | *N. europaea*- 460 $\mu\text{M}$ | 95.4 ± 2.1 (7d) | 3.42±0.2 (7d) | 170.2±1.4 (14d) | 3.42±0.2 (7d) |
| | *N. multiformis*- 460 $\mu\text{M}$ | 36.6±3.4 (8d) | 8.98±0.9 (8d) | 167.6±16.1 (16d) | 10.3±0.8 (14d) |
| **AOA** | “*Ca. N. franklandus*” – 460 $\mu\text{M}$ | 74.9±0.9 (15d) | 8.80±0.1 (15d) | 96.3±1.5 (19d) | 10.3±0.2 (22d) |
| | “*Ca. N. franklandus*” – 46 $\mu\text{M}$ | 4.97±0.7 (15d) | 0.61±0.3 (15d) | 6.75±0.1 (25d) | 0.61±0.4 (22d) |
| | “*Ca. N. franklandus*” – 4.6 $\mu\text{M}$ | 1.78±0.1 (15d) | 0.0 (0.0) | 1.83±0.1 (16d) | 0.0 (0.0) |
| | “*Ca. N. sinensis*” – 460 $\mu\text{M}$ | 132.6±1.5 (7d) | 16.9±0.4 (7d) | 189.8±13.8 (10d) | 27.4±1.1 (17d) |
| | “*Ca. N. sinensis*” – 46 $\mu\text{M}$ | 13.3±0.4 (7d) | 2.3±0.1 (7d) | 13.5±0.7 (6d) | 3.4±1.0 (17d) |
| | “*Ca. N. sinensis*” – 4.6 $\mu\text{M}$ | 1.0±0.0 (7d) | 0.20±0.0 (7d) | 4.3±0.4 (5d) | 0.4±0.1 (27d) |
| **NOB** | *Nitrobacter* sp. - 460 $\mu\text{M}$ | 50.8±13.1 (4d) | 11.5±0.9 (4d) | 114.5±1.8 (9d) | 14.2± 0.6 (3d) |

All other NIs did not show a clear dose-dependent degradation pattern. DMPP and DCD were rather persistent with their DT50 values varying from 4.34 to >1000 days and 45.9 days to >1000 days respectively. In contrast NP was rapidly degraded in most treatments with DT50 values ranging from 0.12 to 12.5 days (Table S1).

## DISCUSSION

We determined *in vitro* the inhibition potency of EQ and its oxidation derivatives, QI and EQNL, on the growth and activity of a range of soil nitrifiers and compared them to the NIs most widely used in agricultural practice (DCD, NP, and DMPP). Five isolates, representative of diverse and globally distributed lineages of soil AO (34, 35) and NOB (36), were tested. *N. europaea* and *N. multiformis* are representatives of AOB clusters 3 and 7, respectively (37), and are commonly used as model soil AOB, with cluster 3 sequences being often the dominant lineage in agricultural and grassland soil ecosystems (38, 39, 34). The two soil AOA isolates represent contrasting ecological niches, with “*Ca*. N. franklandus (40) and “*Ca*. N. sinensis” (41, 42) being representatives of widely distributed neutrophilic and acidophilic AOA lineages, respectively. *Nitrobacter* sp. NHB1 (43) was chosen as a representative of one of the two dominant NOB lineages found in soil (36), with *Nitrobacter* strains typically having greater nitrite oxidation activity compared to those from the genus *Nitrospira* (44, 45).

We first determined the inhibitory potential of EQ to AO and NOB. As a powerful antioxidant, EQ is prone to oxidation, producing a range of transformation products depending on the interacting matrix (e.g. animals, plants, and soils) (46). In this study, EQ was rapidly transformed to QI and EQNL, with the former being the major but least persistent transformation product, while the latter being a minor but more persistent product, in line with previous studies in soil (32, 33). We observed a different inhibition potential for AOB and AOA. All three compounds were more potent inhibitors of the two AOA isolates compared to AOB isolates, with *N. multiformis* being more sensitive than *N. europaea*. EQ was characterized by EC_50_ values lower than those of EQNL, but equal or higher (only in case of *N. multiformis*) than those of QI. Given the transformation pattern of EQ in the microbial cultures, we presume that its calculated EC_50_ values also include the activity of its two oxidative derivatives, QI and EQNL. Considering that in the cultures inhibited by EQ (i) EQNL was formed at concentrations substantially lower than those expected to result in an inhibitory effect on the AO tested, and (ii) QI was formed at concentrations equal or higher than those expected to induce an inhibitory effect on the AO tested, we suggest that QI is the main determinant of the persistent inhibitory effect of EQ on AO and NOB. This is consistent with our previous soil studies where EQ, but mostly QI when applied alone, induced a significant inhibition of potential nitrification and transcription of both bacterial and archaeal *amoA* genes (32). Unlike our previous soil studies where QI showed equivalent inhibitory effects against AOB and AOA, we observed a difference in the inhibition potential of QI between these two AO groups. This discrepancy could be attributed to the concentrations of QI applied or formed in soil samples (up to 86.1 μmol Kg^-1^ dwt soil) which most probably reached the inhibition threshold levels for both AO groups. Further studies under a range of conditions known to affect the activity of AO in soil (e.g. soil pH, NH_4_^+^ amendment) or the performance of NIs (e.g. rate of application, temperature, N source) will determine the potency of these compounds as broad range or AOA-specific NIs.

To establish the full potential of EQ or its derivatives as new potent NIs, we compared them with the *in vitro* inhibitory activity of three established NIs. In contrast to EQ and its derivatives, DCD exhibited higher *in vitro* toxicity towards AOB isolates compared to AOA, in line with most previous soil studies (47–49). In our study, DCD inhibited both AOA isolates at concentrations ≥ 5OO μM. Similar *in vitro* tests with soil enrichment cultures of AOA *Nitrososphaera* sp. JG1 and *Nitrosarchaeum koreense* MY1, showed strong inhibition by DCD at 500 μM (50, 51), while a pure isolate of “*Ca*. Nitrosotalea devanatera” ND1 was inhibited at concentrations >1000 μM (22). Furthermore, Shen *et al.*, (23) reported an EC_50_ of 94O.6 (±85.3) μM for *Nitrososphaera viennensis* EN76. It is interesting to note that “*Ca*. N. sinensis” and “*Ca*. N. franklandus” have different sensitivities to DCD compared to their phylogenetically associated strains “*Ca*. Nitrosotalea devanatera” ND1 and *Nitrososphaera viennensis* EN76, respectively. Despite being closely related, the two *Nitrosotalea* isolates exhibit different physiologies which might explain the observed differences. Although “*Ca*. N. sinensis” (μmax = 0.025 h^-1^) grows approximately twice as fast as “*Ca*. N. devanaterra (ND1)” (μmax = 0.011 h^-1^), their cell yields are similar (4-4.5 cells μM^-1^ NH_3_), while the specific cell activity of ND1 (0.072 fmol NO_2_^-^cell^-1^ h^-1^) is slightly higher than this of “*Ca*. N. sinensis” (0.065 fmol NO_2_^-^cell^-1^ h^-1^) (40). On the other hand, the two *Nitrososphaera* isolates are characterized by similar growth rates (μmax = 0,024 h^-1^). In contrast to our results, a 3x lower EC_50_ was reported by Shen *et al.* (23) for *N. multiformis* (EC_50_= 80.28 (±6.2O) μM vs. 248.7 (±7.4) μM in our study). However, the persistence of DCD in the cultures was not determined.

Similar to DCD, the more recently discovered DMPP showed higher inhibition potency to AOB compared to AOA. For the AOA, “*Ca*. N. franklandus” was less sensitive to DMPP than “*Ca*. N. sinensis”, a difference potentially attributed to the higher specific cell activity of “*Ca*. N. franklandus” (2.02 vs. 0.065 fmol NO_2_^-^cell^-1^ h^-1^ for “*Ca*. N. sinensis”) (40). A differential activity of DMPP towards AOA and AOB has been observed previously in soil studies (28, 52–54), although the bioactivity of DMPP in soil, unlike *in vitro* tests, is influenced by various edaphic, environmental and microbial factors (55). Our study provides the first data on the *in vitro* range activity of DMPP against soil AO.

NP was the only tested NI that showed an equivalent and strong inhibitory effect towards both AOB and AOA isolates, suppressing their activity at concentrations ≥ 1 μM. In accordance with our results, previous studies had found that NP inhibited the activity of *Nitrosomonas, Nitrosospira*, and *Nitrosolobus* strains at concentrations ≥ 0.86 μM (18). Comparative tests with various terrestrial and marine AOA (*Nitrososphaera* sp. JG1, *Nitrosarchaeum koreense* MY1, *Nitrosopumilus maritimus* SCM1, and *Nitrosopumilus cobalaminigenes* HCA1), and AOB (including *N. europaea* and *N. multiformis*), showed an inhibitory effect of NP at 10 μM (56, 51, 57). In line with our findings for “*Ca*. N. sinensis” (EC_50_ = 6.7 μM), Lehtovirta-Morley *et al.*, (22) reported that NP halted the activity and growth of the phylogenetically closely related “*Ca*. N. devanaterra” ND1 at concentrations ≥10 μM. In contrast, Shen *et al.*, (23) reported significantly higher EC_50_ values for NP (> 173 μM and 118.1 μM for *N. multiformis* and *N. viennensis*, respectively). This could be due to the different approach used by Shen *et al.* (23) for adding NP into the cultures. This involved adding non-dissolved NP in the culture medium to achieve concentrations in the range of 40-173 μM, with the highest level corresponding to the upper limit of NP water solubility at 20°C (40 mg L^-1^) entailing a risk for precipitation of the active compound during incubation at 28°C (*N. multiformis*) or 37°C (*N. viennensis*). Soil studies with NP, although limited, are influenced by the relative functional dominance of one group of AO over another and the applied concentration of the inhibitor. Cui *et al.*, (27) reported preferential inhibition of AOB by NP applied at 1.3 μmol Kg^-1^. Conversely, Lehtovirta-Morley *et al.*, (22) found AOA to be sensitive to NP at concentrations ≥10 μmol Kg^-1^ in an acidic soil where “*Ca*. N. devanaterra” ND1 was dominant.

A comparative analysis of the inhibitory range of the tested NIs highlights the serious practical implications of our findings (Fig. 6). DMPP and NP were equally effective and the most potent NIs against AOB, followed by QI, (EQ), and DCD, with QI being more active than DCD against *N. multiformis*. On the other hand, QI (EQ) and NP were equally effective and the most active NIs against AOA, while DCD and DMPP, were not inhibitory to AOA at the concentrations tested (Fig. 6). These findings suggest that NP is the only commercial NI capable of effectively inhibiting both AOB and AOA, and hence the most effective currently available NI, although it is not currently registered for use in Europe. 30% of the World’s soils have a pH <5.5 and European agricultural soils have a mean pH of 5.8 (58). These results therefore have practical implications for low pH soils where ammonia oxidation may be dominated by AOA (41). Slight differences in the inhibition thresholds between AOA and AOB may largely affect agricultural practice, since AOA are expected to contribute to nitrogen fertilizer loss in conditions where AOB would be inhibited (22). On the other hand, universal inhibitory effects on both AOB and AOA (and comammox bacteria) suggest that nitrification inhibition will not be compromised by functional redundancy. Alternatively, the use of mixtures of NIs exhibiting complementary activity against different AO groups or targeting different parts of the ammonia oxidation pathway could be equally efficient with broad range NIs. In this regard, the potential use of EQ as a novel NI is promising, considering its unique feature to be transformed in soil to QI, which is a highly potent inhibitor of AOA, and has a satisfactory inhibitory effect on AOB, comparable with that of established NIs such as DCD.

**Fig. 6.**
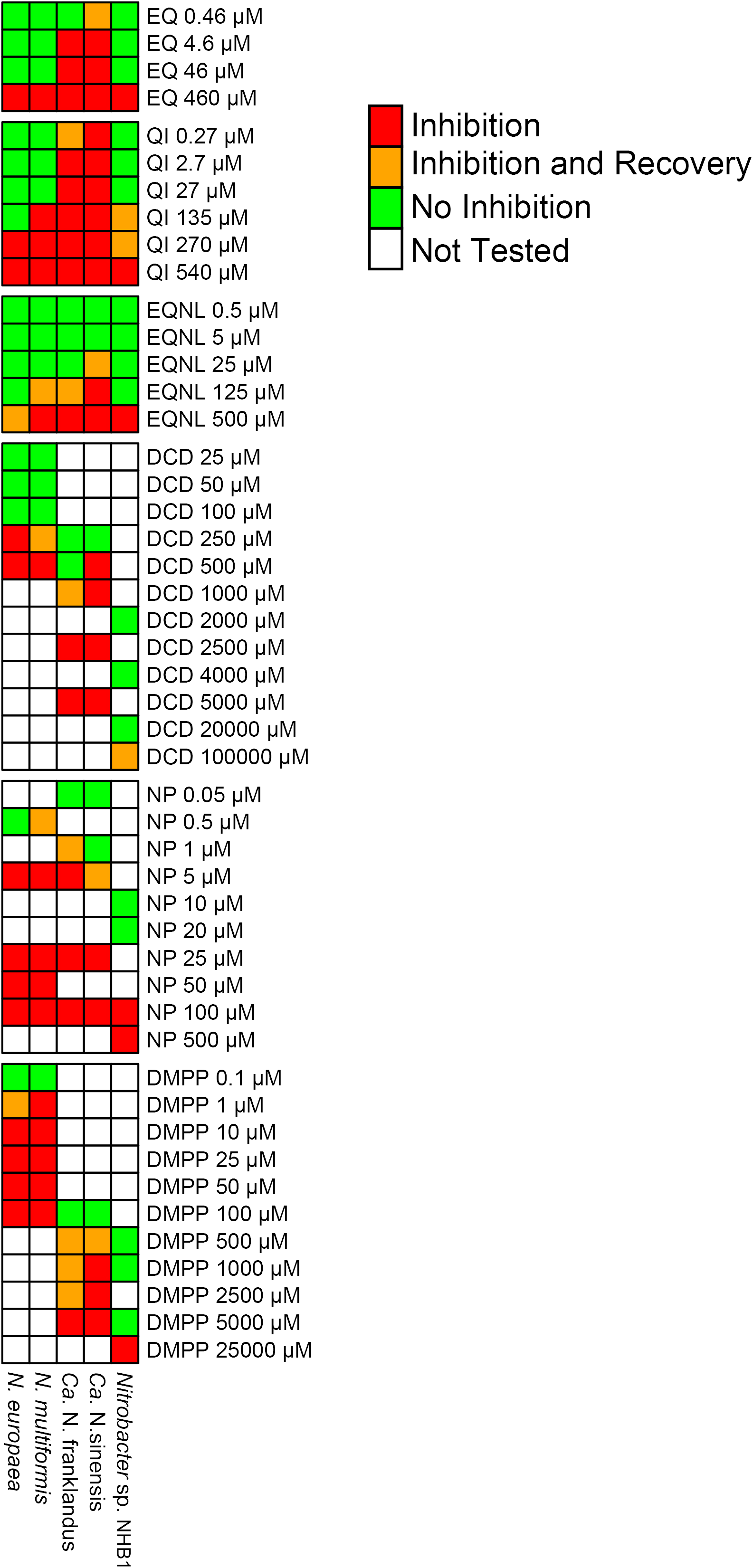
Heatmap representation of the qualitative impact of different concentrations of NIs on the nitrifying activity of soil ammonia- and nitrite-oxidizing isolates. The level of inhibition observed was classified in the following categories: “Inhibition” (shown in red); “Inhibition and Recovery” (shown in orange); “No inhibition” (shown in green); “Not tested” (shown in white).

In addition to the practical implications, the considerable differences in the range of inhibitory activities exhibited by the tested NIs might indicate differences in their mode of action not considered before. For example, DCD, DMPP and NP, all considered as Cu-chelators, showed variable activity towards AOA, which, unlike AOB, rely on copper-containing proteins for electron transfer (59). NP on the other hand was previously proposed to serve also as an alternative AMO substrate, generating products (6-chloropicolinic acid) that irreversibly deactivate ammonia oxidation (10). This dual inhibitory mechanism of NP might offer an explanation for its more universal inhibitory activity towards both AOA and AOB. EQ and its derivatives possess high-antioxidative capacity acting as free radical scavengers (46). As EQ and QI showed similar inhibitory effects to other NO-scavengers (e.g. PTIO) (57), their effectivity against AOA may be due to a similar mode of action. Alternatively, as QI is a strong antioxidant, it could be involved in oxidative stress-related cell disruption particularly in AOA, with AOB being capable of coping with oxidative stress using catalases which are largely absent in AOA (60). Further studies should define the inhibition mechanism of EQ on AO and clarify the corresponding mechanisms of the other NIs which remain unknown.

Given the effect of NIs on AO, they will also have an indirect inhibitory effect on NOB, the functional partners of AO. We demonstrated that DMPP and DCD, despite their strong impact on AOB, had no inhibitory effect on *Nitrobacter* sp. NHB1, in contrast to NP, EQ and its derivatives which were more active against *Nitrobacter* sp. NHB1. Previous soil studies also demonstrated that DCD was not suppressive to *Nitrospira*- and *Nitrobacter-like* bacteria at levels up to 150 μmol Kg^-1^ soil (61, 62, 49). To date there are no data, either *in vitro* or in soil, regarding the impact of DMPP or EQ and its derivatives on NOB, while NP applied at rates up to 50 μM did not inhibit the nitrite-oxidizing activity of the widely distributed *Nitrobacter agilis* (20). We provide the first evidence for the toxicity of NIs on a *Nitrobacter* sp. Further studies extended to other NOB, including the widely distributed and diverse *Nitrospira*-like bacteria, would determine the full inhibitory potential of NIs on soil NOB.

In parallel to activity and growth measurements, we determined the degradation and transformation of the tested NIs to identify potential links between the duration of exposure (persistence) and the effects observed. The total residues of EQ showed limited persistence in the AOA and NOB cultures (DT_50_ = 2.4-8.7 days), and low to moderate persistence in the AOB cultures (DT_50_ = 8.7-60.1 days), a difference most likely attributed to abiotic factors such as medium pH (acidic for AOA and NOB vs. alkaline for AOB) rather than an enzymatic transformation, considering the autotrophic lifestyle of the tested isolates (60) and the recalcitrance of EQ under aerobic and anaerobic conditions (63). However, a direct interaction of these compounds with the tested organisms cannot not be excluded (10). The three commercial NIs showed remarkably different stability in the liquid cultures. DCD showed moderate to high persistence (DT_50s_ = 44.5 to >1000 days), whereas NP degraded rapidly (DT_50_ =0.12-12.5 days). DMPP showed a high persistence in all liquid cultures, with the exception of *Nitrobacter* cultures where a great variation in the persistence of DMPP was observed. Considering that *Nitrobacter* sp. NHB1 and AOA were cultured in media of similar content and pH, the above variation was possibly induced by interaction between the NI and the *Nitrobacter* sp. strain. Overall, we did not observe any clear correlations between NIs persistence and inhibition potency.

## Conclusions

We determined *in vitro* the inhibition potential of novel (EQ and its derivatives) and all NIs currently used in the agricultural practice on representative soil-derived AOA, AOB, and NOB. EQ, and primarily its major transformation product QI, showed high potency against AOA, in contrast to DCD and DMPP, the only NIs currently registered for use in Europe and which are inhibitory only to AOB. In contrast, NP showed an inhibitory activity against all groups tested. The activity of those NIs on comammox bacteria are still unknown due to the lack of soil-derived isolates, and their characterization will be required to provide a complete understanding of their inhibition potency on all AO.

Our study (i) offers benchmarking knowledge of the activity range of known and potentially new NIs to soil AO and *Nitrobacter* NOB, (ii) introduces the novel potential NI EQ, which possesses desirable characteristics including transformation into another potent NI, QI, which has high potency against AOA in contrast to other registered NIs in Europe, and (iii) demonstrates the different sensitivity of AOA and AOB to NIs, providing potentially novel strategies relying on new broad-range NIs, or more likely, using mixtures of NIs which possess complementary activity against different nitrifier groups. Further elucidation of EQ and QI inhibitory mechanisms, and verification of their efficacy under diverse soil conditions affecting both the activity of AO and the performance of NIs, might lead to the development of a novel NI for more efficient N conservation in agricultural soils.

## MATERIAL AND METHODS

### Microbial strains, growth conditions and chemicals

Five soil-derived nitrifying isolates were used in the *in vitro* assays: two AOB (*N. europaea, N. multiformis*), two AOA (“*Ca*. N. franklandus”, “*Ca*. N. sinensis”), and one NOB (*Nitrobacter* sp. NHB1). All strains were grown aerobically in the dark without shaking. Details on the cultivation media and incubation temperatures used are given in the Supplemental Material.

Analytical standards of DCD (99 % purity), NP (≥ 98 %), and EQ (95%) were purchased from Sigma-Aldrich (Germany), while the DMPP (99.1%) analytical standard was provided by BASF Hellas. The oxidation derivatives of EQ, QI and EQNL were synthesized as described by Thorisson *et al.* (64). The chemical structures of all studied compounds are shown in Fig. S7.

### Liquid culture assays

The activity of all NIs was determined in liquid batch cultures over a broad range of concentrations to establish relevant inhibition thresholds (EC_50_ values) per strain and compound. For each, triplicate strain x NI x concentration replicates were established in 100-mL Duran bottles containing 50 mL of growth medium and inoculated with a 1 or 2% (v/v) transfer from exponentially growing cultures of AOB or AOA/NOB, respectively. EQ, QI, EQNL and NP were added to the cultures as filter sterilized DMSO solutions due to their low water solubility (≤60 mg L^-1^ at 2O°C) and the final concentration of DMSO in all cultures was 0.1% (v/v). DCD and DMPP were dissolved in sterile dH_2_O before addition of 25 μl (0.5% v/v) in the different cultures. All NIs were added to batch cultures at the beginning of the exponential growth phase. For all assays, cultures were established in triplicate with the same inoculum without NI amendment. Upon inoculation all liquid batch cultures were sampled at regular time intervals to determine the effect of NIs on the activity and growth of AO by measuring changes in nitrite concentrations, and *amoA* or *nxrB* gene abundance for AO and NOB populations, respectively.

### Nitrite measurements and gene abundance quantification

Nitrite concentrations were determined colorimetrically at 540 nm in a 96-well plate format assay by diazotizing and coupling with Griess reagent (65). *amoA* and *nxrB* gene abundance was determined in a Biorad CFX Real-Time PCR system. DNA was extracted from a cell pellet obtained from 2-ml aliquots of the microbial cultures using the tissue DNA extraction kit (Macherey-Nagel, Germany). The *amoA* genes of AOB and AOA was amplified with primers amoA-1F/amoA-2R (66) and Arch-amoAF/Arch-amoAR (67), respectively as described by Rousidou *et al.*, (68). The *nxrB* gene of *Nitrobacter* was quantified with primers nxrB-1F and nxrB-1R (69) using the following thermal cycling conditions: 95°C for 3 min, followed by 40 cycles of 95°C for 30 seconds, 57°C for 20 seconds, 72°C for 30 seconds, with a final dissociation curve analysis. The abundance of *amoA* and *nxrB* genes were determined via external standard curves as described by Rousidou *et al.*, (68). qPCR amplification efficiencies ranged from 80.3% to 109.4%, with *r*^2^ values ≥ 0.98.

### Nitrification Inhibitors extraction

EQ, QI, EQNL, and NP residues were extracted from liquid media by mixing 0.3 mL liquid culture with 0.7 mL of acetonitrile. Residues of DCD and DMMP were extracted by mixing 0.1 mL liquid culture with 0.9 mL of ddH_2_O water and methanol, respectively. The derived mixtures were vortexed for 30 s and stored at −20°C until analysis. Recovery tests at three concentration levels (in the range of the tested concentrations) showed recoveries of >80% for all compounds studied.

### Chromatographic analyses

High performance liquid chromatography (HPLC) analyses were performed in a Shimadzu LC-20ADHPLC system equipped with an UV/VIS PDA detector. A Shimadzu GVP-ODs (4.6 mm by 150mm, 5μm) pre-column, connected to a RP Shimadzu VP-ODs (4.6 mm x 150 mm, 5μm) column, was used for NI separation. The injection volume was 20 μl. The flow rate of the mobile phase was set at 0.8 mL min^-1^ for DCD, and at 1 mL min^-1^ for all other NIs. Column temperature was set at 40°C for DCD and DMPP, and at 25 °C for all the other NIs. Mixtures of acetonitrile and ammonia (0.25% [vol/vol]) or *ortho*-phosphoric acid (0.1% [vol/vol]) were used at a ratio of 70:30 (vol/vol) for mobile phases in the analyses of EQ, QI, EQNL, and NP, respectively, and detection was achieved at 225, 245, 230, and 269 nm, respectively. Similarly, chromatographic separation of DCD and DMPP was achieved using ddH_2_O (100%) and a mixture of methanol and *ortho*-phosphoric acid (0.1% [vol/vol]) solution 50:50 by volume, respectively. DCD and DMPP residues were detected at 218 nm and 225 nm, respectively.

### Calculation of inhibition threshold levels (EC_50_)

In this study, EC_50_ describes the concentration of the inhibitor that reduces half of the activity (nitrite accumulation or consumption) of AO or NOB, with dose-response modeling performed using normalized data whereby nitrite concentration values were divided by the mean value of the matching control. Analyses were carried out using the dose response curves (drc) v3.0-1 packagse (70) of the R software (71). A brief description of the tested models can be found in Ritz *et al.*, (72). An empirical modeling approach was initially used for selecting the best fitting model according to tested goodness of fit indices (see Supplemental Material), followed by the choice of the four-parameter log logistic model as the best compromise among tested models for comparing endpoint values.

### Data analysis

Nitrite and qPCR data were subjected to one-way ANOVA, followed by Tukey’s post hoc test (*P*<0.05). Variance between the EC_50_ values of the different NIs for one strain and between different strains for a given NI was analyzed by one-way ANOVA, and Duncan post hoc test (P < 0.05). The four kinetic models proposed by the FOCUS working group on pesticide degradation kinetics (73) (SFO and the biphasic models hockey stick (HS), first order multi-compartment (FOMC), and double first order in parallel (DFOP)) were used to calculate NI degradation kinetic parameters (DT_50_, k_deg_). Curve fitting was performed with the mkin v0.9.47.1 (74) package of the R v3.4.3 software (71). More details on degradation kinetics are given in the Supplemental Material.

## ACKNOWLEDGMENTS

This work is part of the project “NITRIC - Looking up for **N**ovel n**ITR**ification **I**nhibitors: New stories with old **C**ompounds” which has received funding from the Hellenic Foundation for Research and Innovation (HFRI) and the General Secretariat for Research and Technology (GSRT), under grant agreement No. 1229.

The authors declare no conflict of interest.

## Abbreviations

NIs: nitrification inhibitors
EQ: ethoxyquin
QI: 2,6-dihydro-2,2,4-trimethyl-6-quinone imine
EQNL: 2,4-dimethyl-6-ethoxyquinoline
DCD: dicyandiamide
NP: nitrapyrin
DMPP: 3,4-dimethylpyrazole phosphate
AOB: ammonia-oxidizing bacteria
AOA: ammonia-oxidizing archaea
AO: ammonia-oxidizers
NOB: nitrite-oxidizing bacteria
comammox: complete ammonia-oxidizing bacteria
AMO: ammonia monooxygenase

