## Supplemental Material for "Comparison of the *in vitro* activity of novel and established nitrification inhibitors applied in agriculture: challenging the effectiveness of the currently available compounds"

Dr. Evangelia S. Papadopoulou

**Abbreviations[1]**

[1]NIs: nitrification inhibitors, EQ: ethoxyquin; QI: 2,6-dihydro-2,2,4-trimethyl-6-quinone imine; EQNL: 2,4-dimethyl-6-ethoxyquinoline; DCD: dicyandiamide, NP: nitrapyrin, DMPP: 3,4-dimethylpyrazole phosphate, AOB: ammonia-oxidizing bacteria, AOA: ammonia-oxidizing archaea, AO: ammonia-oxidizers, NOB: nitrite-oxidizing bacteria, comammox: complete ammonia-oxidizing bacteria; AMO: ammonia monooxygenase

### MATERIALS AND METHODS

**Microbial strains, growth conditions and chemicals.** Five soil-derived nitrifying isolates were used in the *in vitro* assays: two AOB (*N. europaea*, *N. multiformis*), two AOA ("*Ca.* N. franklandus", "*Ca.* N. sinensis"), and one NOB (*Nitrobacter* sp. NHB1). All strains were grown aerobically in the dark without shaking. "*Ca*. Nitrosocosmicus franklandus" was isolated from a Scottish agricultural soil with pH 7.5 (1), and "*Ca*. Nitrosotalea sinensis" was isolated from a Chinese acidic paddy soil (pH 4.7) (2). AOB were grown at 28°C in Skinner and Walker's medium (3) (pH of 7.5). AOA were incubated at 35°C in a medium supplemented with 1 mM $NH_4^+$ ($NH_4Cl$). "*Ca*. Nitrosocosmicus franklandus" was cultured in HEPES-buffered modified fresh water medium (pH 7.5) (1), while "*Ca*. Nitrosotalea sinensis" was grown in a freshwater medium (pH 5.2) (4). *Nitrobacter* sp. (5) was grown at 28°C in fresh water medium (pH 5.2) (4) supplemented with 0.5 mM $NO_2^-$ ($NaNO_2$).

Analytical standards of DCD (99 % purity), NP (≥ 98 %), and EQ (95%) were purchased from Sigma-Aldrich (Germany), while the DMPP (99.1%) analytical standard was provided by BASF Hellas. The oxidation derivatives of EQ, QI and EQNL were synthesized as described by Thorisson *et al*. (6). The chemical structures of all studied compounds are shown in Fig. S7.

**Liquid culture assays.** The activity of all NIs was determined in liquid batch cultures over a broad range of concentrations to establish relevant inhibition thresholds ($EC_{50}$ values) per strain and compound. For each, triplicate strain x NI x concentration replicates were established in 100-mL Duran bottles containing 50 mL of growth medium and inoculated with a 1 or 2% (v/v) transfer from exponentially growing cultures of AOB or AOA/NOB cultures, respectively. EQ, QI, EQNL and NP were added to the cultures as filter sterilized DMSO solutions due to their low water solubility (≤60 mg $L^{-1}$ at 20°C) and the final concentration of DMSO in all cultures was 0.1% (v/v). DCD and DMPP were dissolved in sterile $dH_2O$ before addition of 25 μl (0.5% v/v). All NIs were added to batch cultures at the beginning of the exponential growth phase. For all assays, cultures were established in triplicate with the same inoculum without NI amendment. Upon inoculation all liquid batch cultures were sampled at regular time intervals to determine the effect of NIs on the activity and growth by measuring changes in nitrite concentrations, and *amoA* or *nxrB* gene abundance for AO and NOB populations, respectively.

**Nitrite measurements and gene abundance quantification.** Nitrite concentrations were determined colorimetrically in a 96-well plate format assay by diazotizing and coupling with Griess reagent (7). For each measurement duplicate standards ranged between 0 and 100 μM $NaNO_2$ were used. Absorbance was recorded at 540 nm, using an Enspire multimode plate reader (PerkinElmer). *amoA* and *nxrB* gene abundance was determined in a Biorad CFX Real–Time PCR system. DNA was extracted from a cell pellet obtained from 2-ml aliquots of the microbial cultures using the tissue DNA extraction kit (Macherey-Nagel, Germany). The *amoA* gene of AOB and AOA was amplified with primers amoA-1F/amoA-2R (8) and Arch-amoAF/Arch-amoAR (9), respectively. The *nxrB* gene of *Nitrobacter* was quantified with primers nxrB-1F and nxrB-1R (10). Amplifications were performed using KAPA SYBR Fast universal 2x qPCR master mix according to the manufacturer's instructions. The thermal cycling conditions used for AO were as described elsewhere (10), while for NOB the following thermal cycling conditions were used: 95°C for 3 mins, followed by 40 cycles of 95°C for 30 seconds, 57°C for 20 seconds, 72°C for 30 seconds, with a final dissociation curve analysis. The copy numbers of the *amoA* and *nxrB* genes were determined via external standard curves as described by Rousidou *et al*., (11). qPCR amplification efficiencies ranged from 80.3% to 109.4%, with $r^2$ values ≥ 0.98.

**Nitrification Inhibitors extraction.** EQ, QI, EQNL, and NP residues were extracted from liquid media by mixing 0.3 mL liquid culture with 0.7 mL of acetonitrile. Residues of DCD and DMPP were extracted by mixing 0.1 mL liquid culture with 0.9 mL of $ddH_2O$ and methanol, respectively. The derived mixtures were vortexed for 30 s and stored at −20˚C until analysis. Recovery tests at three concentration levels (in the range of the tested concentrations) showed recoveries of >80% for all compounds studied.

**Chromatographic analyses.** High performance liquid chromatography (HPLC) analyses were performed in a Shimadzu LC-20ADHPLC system equipped with an UV/VIS PDA detector. A Shimadzu GVP-ODs (4.6 mm by 150mm, 5μm) pre-column, connected to a RP Shimadzu VP-ODs (4.6 mm x 150 mm, 5μm) column, was used for NI separation. The injection volume was 20 μl. The flow rate of the mobile phase was set at 0.8 mL $min^{-1}$ for DCD and at 1 mL $min^{-1}$ for all the other NIs. Column temperature was set at 40˚C for DCD and DMPP, and at 25˚C for all the other compounds. Mixtures of acetonitrile and ammonia (0.25% [vol/vol]) or *ortho*-

phosphoric acid (0.1% [vol/vol]) were used at a ratio of 70:30 (vol/vol) for mobile phases in the analyses of EQ, QI, EQNL, and NP, respectively, and detection was achieved at 225, 245, 230, and 269 nm, respectively. Similarly, chromatographic separation of DCD and DMPP was achieved using $ddH_2O$ (100%) and a mixture of methanol and *ortho*-phosphoric acid (0.1% [vol/vol]) solution 50:50 by volume, respectively. DCD and DMPP residues were detected at 218 nm and 225 nm, respectively.

**Calculation of inhibition threshold levels ($EC_{50}$).** In this study, $EC_{50}$ describes the concentration of the inhibitor that reduces half of the activity (nitrite accumulation or consumption) of AO or NOB, with dose-response modeling performed using normalized data whereby nitrite concentration values were divided by the mean value of the matching control. Analyses were carried out using the dose response curves (drc) v3.0-1 package (12) of the R software (13). A brief description of the tested models can be found in Ritz *et al,* (14). An empirical modelling approach was used for selecting the best fitting model. Goodness of fit (GoF) indices used in the model selection were: Akaike information criterion (AIC) values (used as the major criterion for model assessment); standard deviation of the residuals; standard error of the $EC_{50}$ values; weighted coefficient of determination ($R^2$) as implemented in the qpcR v1.4-0 package (15) (based on the models lack of linearity as opposed to the standard coefficient of determination (16); and the lack of fit approach. The four-parameter log logistic model was used as the best compromise between models in order to make more valid comparisons.

**Data analysis.** Nitrite and qPCR data were subjected to one-way analysis of variance (ANOVA), followed by Tukey's post hoc test ($P<0.05$) to identify significant effects of the NIs at each time point. Variance between the $EC_{50}$ values of the different NIs within a microorganism and between different microorganisms for a given inhibitor was analyzed using one-way ANOVA and following Duncan post hoc test ($P < 0.05$). The four kinetic models proposed by the FOCUS working group on pesticide degradation kinetics (17) were used to calculate NIs degradation kinetic parameters ($DT_{50}$, $k_{deg}$): the single first order kinetic model (SFO), and the biphasic models hockey stick (HS), first order multi-compartment model (FOMC) and double first order in parallel model (DFOP). The goodness of fit was assessed using the $\chi^2$ test as well as visual inspection and the distribution of the residuals. In general, the biphasic kinetic models were used only in cases where the SFO model failed to acceptably describe ($\chi^2$> 15%) inhibitors dissipation. For the dissipation kinetics of the transformation products, the guidelines of the FOCUS working group on pesticides degradation kinetics were followed (17). Curve fitting was performed with the mkin v0.9.47.1 (18) package of the R v3.4.3 software (12).

**TABLE S1.** The $DT_{50}$ values (days) of the different nitrification inhibitors (NIs) tested per nitrifying microorganism and NI concentration tested. $DT_{50}$ values were calculated by fitting the degradation data to the best fitting kinetic model as described in the Experimental Procedures. The first order kinetic model provided the best fit to the experimental data in most cases. In all other cases the model providing the best fit is indicated (i.e. HS: Hockey-Stick model; FOMC: First order multicompartment model; DFOP: Double first order in parallel model)

| Nitrification Inhibitor | Concentration (µM) | *Nitrosomonas europaea* | *Nitrosospira multiformis* | "*Ca.* N. franklandus" | "*Ca.* N. sinensis" | *Nitrobacter sp.* NHB1 |
|---|---|---|---|---|---|---|
| **Total Ethoxyquine (EQ) residues** | **46** | 8.68 | 11.6 | 4.96 | 2.25 | 3.66 |
| | **460** | 48.5 | 60.1 | 8.72 | 2.42 | 2.06 |
| **Quinone Imine (QI)** | **2.7** | 0.05 | 0.52 | 1.39 | 1.52 | 0,90 |
| | **27** | 0.42 | 0.92 | 1.26 | 0.85 | 0,54 |
| | **135** | 1.19 | 1.32 | 1.74 | 1.38 | 0.92 |
| | **270** | 1.91 | 2.47 | 2.68 | 1.92 | 1.18 |
| | **540** | 3.55 | 5.65 | 4.50 | 2.93 | 2.23 |
| **Ethoxyquinoline (EQNL)** | **5** | 7.31 | >1000 | 290.8 | 699.9 | 123.5 |
| | **25** | 10.92 | 18.3 | 289.8 | >1000 | 119.6 |
| | **125** | 12.16 | 73.9 | 400 | >1000 | >1000 |
| | **500** | 82.5 | >1000 | 86.9 | >1000 | >1000 |
| **Dicyanamide (DCD)** | **25** | 66.9 | 111.9 | | | |
| | **50** | 402.3 | >1000 | | | |
| | **100** | >1000 | 109.6 | | | |
| | **250** | 106.4 | 78.5 | >1000 | 225.9 | |
| | **500** | 78.3 | 55.5 | 494.1 | 356 | |
| | **1000** | | | >1000 | >1000 | |
| | **2000** | | | | | 46.9 |
| | **2500** | | | 997.5 | >1000 | |
| | **4000** | | | | | 47.8 |

| | | | | | | |
|---|---|---|---|---|---|---|
| | **5000** | | | >1000 | >1000 | |
| | **20000** | | | | | 45.9 |
| | **100000** | | | | | 84.0 |
| **Nitrapyrin (NP)** | **1** | | | 1.43 | 1.28 | |
| | **5** | 1.53 | $0.82^{HS}$ | $0.12^{FOMC}$ | 1.90 | |
| | **10** | | | | | 2.43 |
| | **20** | | | | | 3.24 |
| | **25** | 2.85 | $0.76^{HS}$ | $0.13^{DFOP}$ | 1.85 | |
| | **50** | 3.40 | $0.56^{FOMC}$ | | | |
| | **100** | 3.24 | 4.01 | $0.15^{DFOP}$ | 2.10 | 4.02 |
| | **500** | | | | | 12.5 |
| **DMPP** | **1** | >1000 | >1000 | | | |
| | **10** | >1000 | >1000 | | | |
| | **25** | >1000 | >1000 | | | |
| | **50** | >1000 | >1000 | | | |
| | **100** | >1000 | >1000 | >1000 | >1000 | |
| | **500** | | | >1000 | >1000 | 46.9 |
| | **1000** | | | >1000 | >1000 | 37.8 |
| | **2500** | | | >1000 | >1000 | |
| | **5000** | | | >1000 | >1000 | >1000 |
| | **25000** | | | | | 4.34 |

### ACKNOWLEDGMENTS

This work is part of the project "NITRIC – Looking up for **N**ovel n**ITR**ification **I**nhibitors: New stories with old **C**ompounds" which has received funding from the Hellenic Foundation for Research and Innovation (HFRI) and the General Secretariat for Research and Technology (GSRT), under grant agreement No. 1229.

### SUPPLEMENTARY FIGURE LEGENDS

**FIG. S1.** The effect of different concentrations of NIs on the growth of *Nitrosomonas europaea* assessed by q-PCR of *amoA* gene abundance. Within each time point bars designated by different lower-case letters are significantly different at the 5% level. Error bars represent the standard error of the mean of biological triplicates.

**FIG. S2.** The effect of different concentrations of the NIs on the growth of *Nitrosospira multiformis* assessed by q-PCR of *amoA* gene abundance. Within each time point bars designated by different lower-case letters are significantly different at the 5% level. Error bars represent the standard error of the mean of biological triplicates.

**FIG. S3.** The effect of different concentrations of the NIs on the growth of "*Candidatus* Nitrosocosmicus franklandus" assessed by q-PCR of *amoA* gene abundance. Within each time point bars designated by different lower-case letters are significantly different at the 5% level. Error bars represent the standard error of the mean of biological triplicates.

**FIG. S4.** The effect of different concentrations of the NIs on the growth of "*Candidatus* Nitrosotalea sinensis" assessed by q-PCR of *amoA* gene abundance. Within each time point bars designated by different lower-case letters are significantly different at the 5% level. Error bars represent the standard error of the mean of biological triplicates.

**FIG. S5.** The effect of different concentrations of the NIs on the growth of *Nitrobacter* sp., assessed by q-PCR of the *nxrB* gene. Within each time point bars designated by different lower-case letters are significantly different at the 5% level. Error bars represent the standard error of the mean of biological triplicates.

**FIG. S6.** The transformation of different concentration levels of Ethoxyquin (EQ) in the nitrifying strains cultures. Each value is the mean of triplicates ± standard error. Bars designated by one asterisk show the concentrations (μM) of EQ and its oxidative derivatives at the onset of inhibition, while bars designated by two asterisks indicate the time point when maximum QI concentrations (μM) were observed.

**FIG. S7.** The chemical structures of the tested nitrification inhibitors (NIs)

**FIG. S8.** Degradation of Quinone Imine in the liquid cultures of the nitrifying isolates tested.

**FIG. S9.** Degradation of Ethoxyquinoline in the liquid cultures of the nitrifying isolates tested.

**FIG. S10.** Degradation of DCD in the liquid cultures of the nitrifying isolates tested.

**FIG. S11.** Degradation of Nitrapyrin in the liquid cultures of the nitrifying isolates tested.

**FIG. S12.** Degradation of DMPP in the liquid cultures of the nitrifying isolates tested.

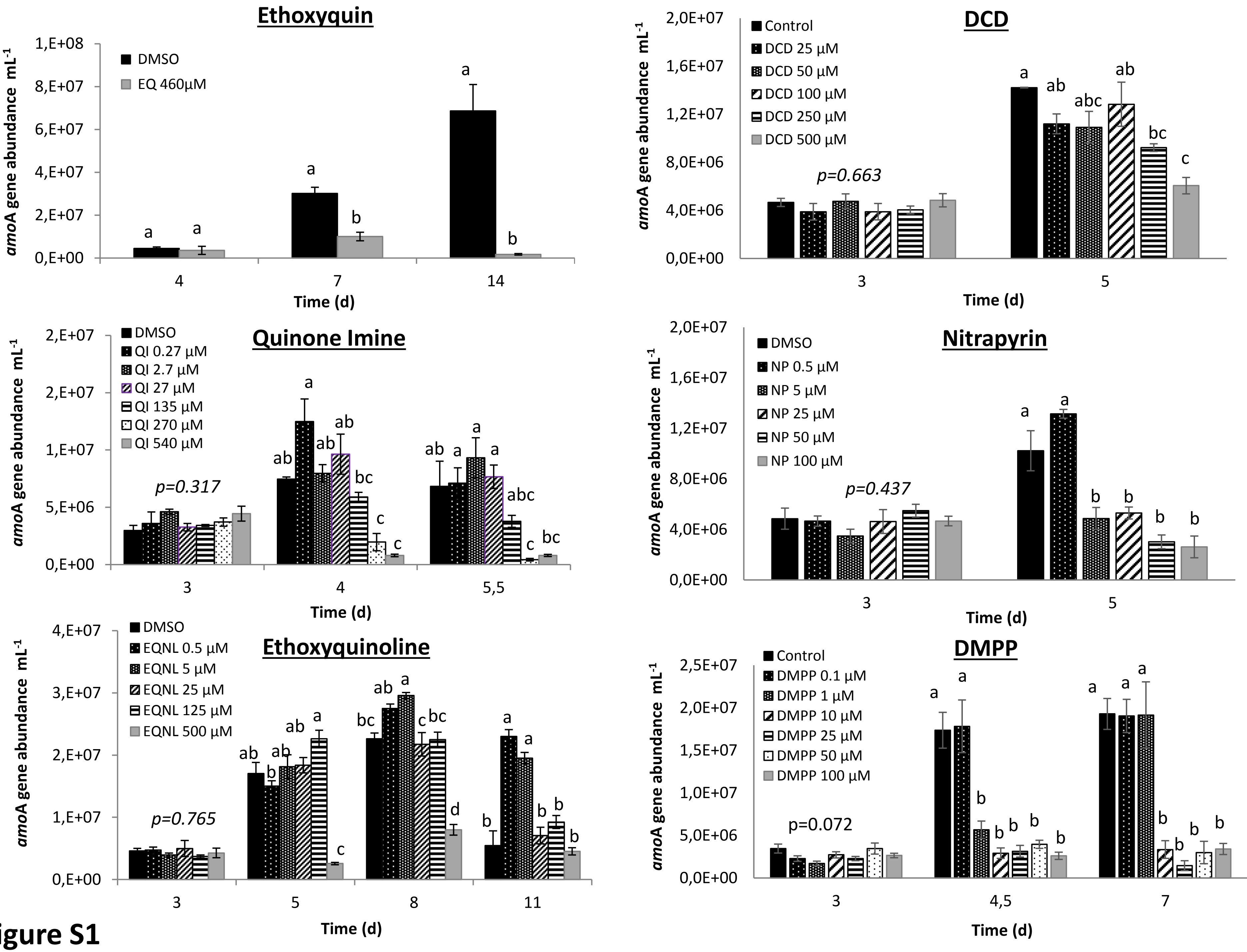

Ethoxyquin
DMSO
EQ 460µM
amoA gene abundance mL-1
Time (d)
DCD
Control
DCD 25 µM
DCD 50 µM
DCD 100 µM
DCD 250 µM
DCD 500 µM
p=0.663
Quinone Imine
DMSO
QI 0.27 µM
QI 2.7 µM
QI 27 µM
QI 135 µM
QI 270 µM
QI 540 µM
p=0.317
Nitrapyrin
DMSO
NP 0.5 µM
NP 5 µM
NP 25 µM
NP 50 µM
NP 100 µM
p=0.437
Ethoxyquinoline
DMSO
EQNL 0.5 µM
EQNL 5 µM
EQNL 25 µM
EQNL 125 µM
EQNL 500 µM
p=0.765
DMPP
Control
DMPP 0.1 µM
DMPP 1 µM
DMPP 10 µM
DMPP 25 µM
DMPP 50 µM
DMPP 100 µM
p=0.072


**Figure S1**

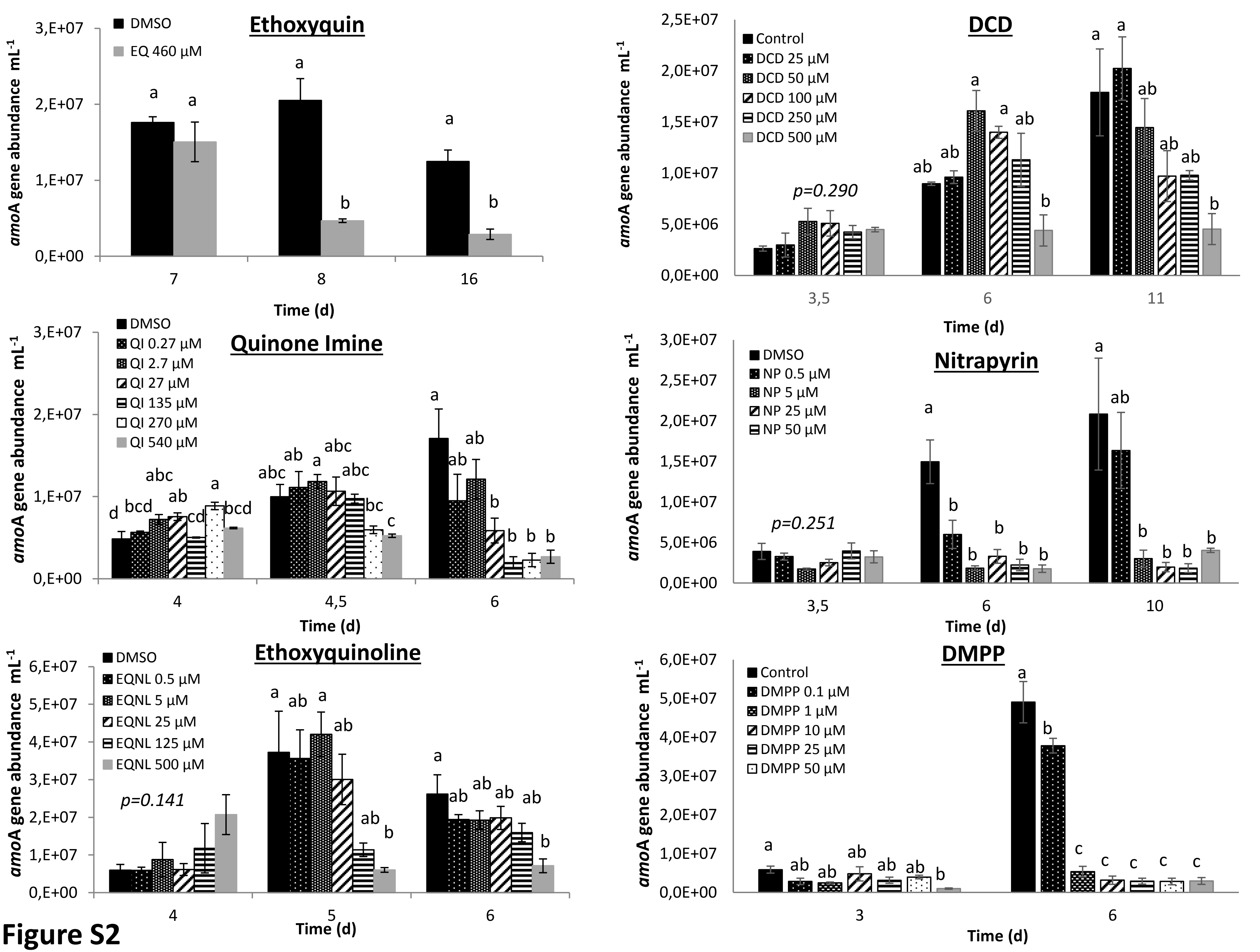

Ethoxyquin
DMSO
EQ 460 µM
amoA gene abundance mL-1
Time (d)
DCD
Control
DCD 25 µM
DCD 50 µM
DCD 100 µM
DCD 250 µM
DCD 500 µM
p=0.290
Quinone Imine
QI 0.27 µM
QI 2.7 µM
QI 27 µM
QI 135 µM
QI 270 µM
QI 540 µM
Nitrapyrin
NP 0.5 µM
NP 5 µM
NP 25 µM
NP 50 µM
p=0.251
Ethoxyquinoline
EQNL 0.5 µM
EQNL 5 µM
EQNL 25 µM
EQNL 125 µM
EQNL 500 µM
p=0.141
DMPP
DMPP 0.1 µM
DMPP 1 µM
DMPP 10 µM
DMPP 25 µM
DMPP 50 µM


**Figure S2**

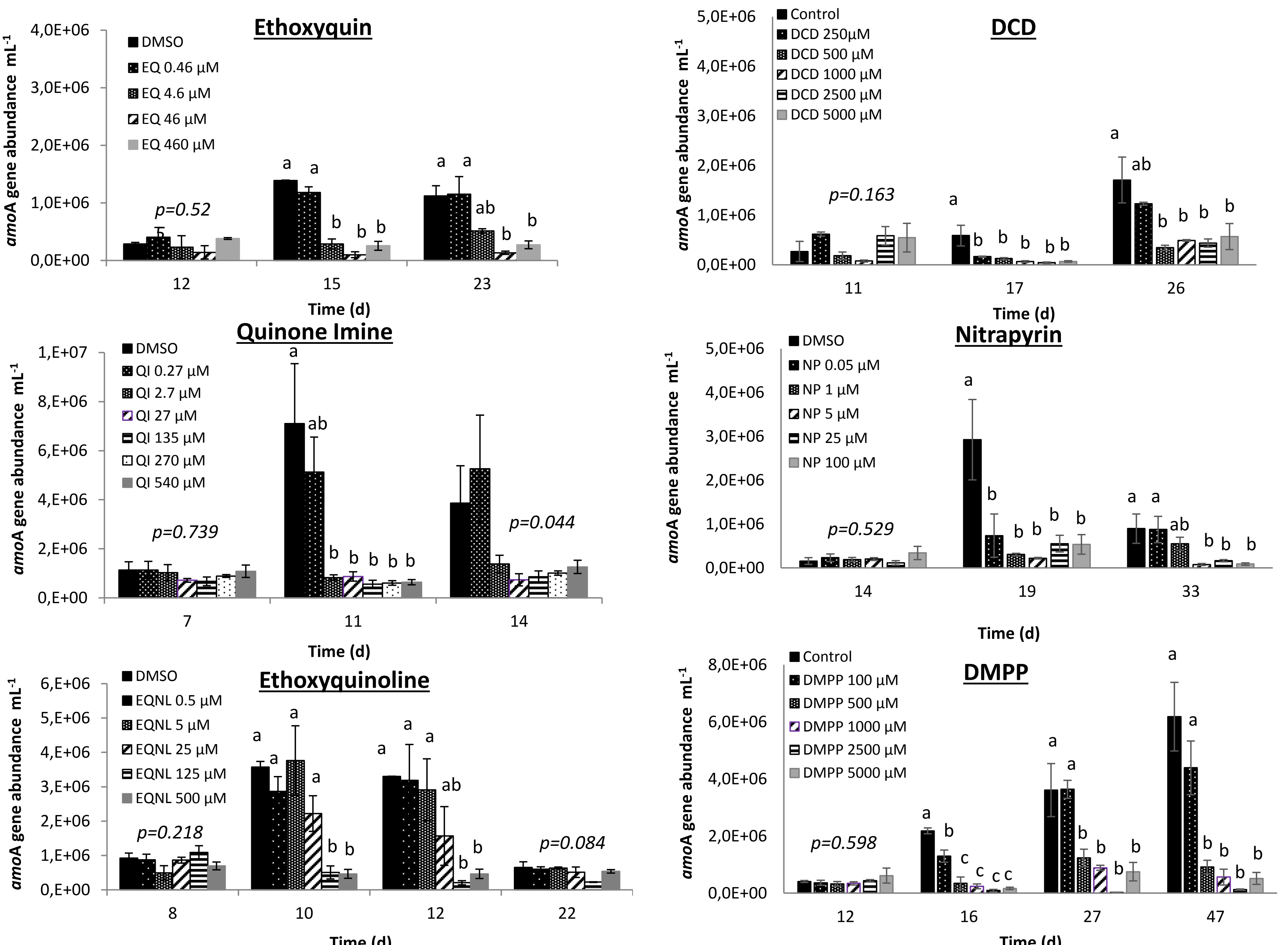
Ethoxyquin
DMSO
EQ 0.46 µM
EQ 4.6 µM
EQ 46 µM
EQ 460 µM
p=0.52
amoA gene abundance mL-1
Time (d)
12
15
23
DCD
Control
DCD 250µM
DCD 500 µM
DCD 1000 µM
DCD 2500 µM
DCD 5000 µM
p=0.163
11
17
26
Quinone Imine
QI 0.27 µM
QI 2.7 µM
QI 27 µM
QI 135 µM
QI 270 µM
QI 540 µM
p=0.739
p=0.044
7
11
14
Nitrapyrin
NP 0.05 µM
NP 1 µM
NP 5 µM
NP 25 µM
NP 100 µM
p=0.529
14
19
33
Ethoxyquinoline
EQNL 0.5 µM
EQNL 5 µM
EQNL 25 µM
EQNL 125 µM
EQNL 500 µM
p=0.218
p=0.084
8
10
12
22
DMPP
DMPP 100 µM
DMPP 500 µM
DMPP 1000 µM
DMPP 2500 µM
DMPP 5000 µM
p=0.598
12
16
27
47

**Figure S3**

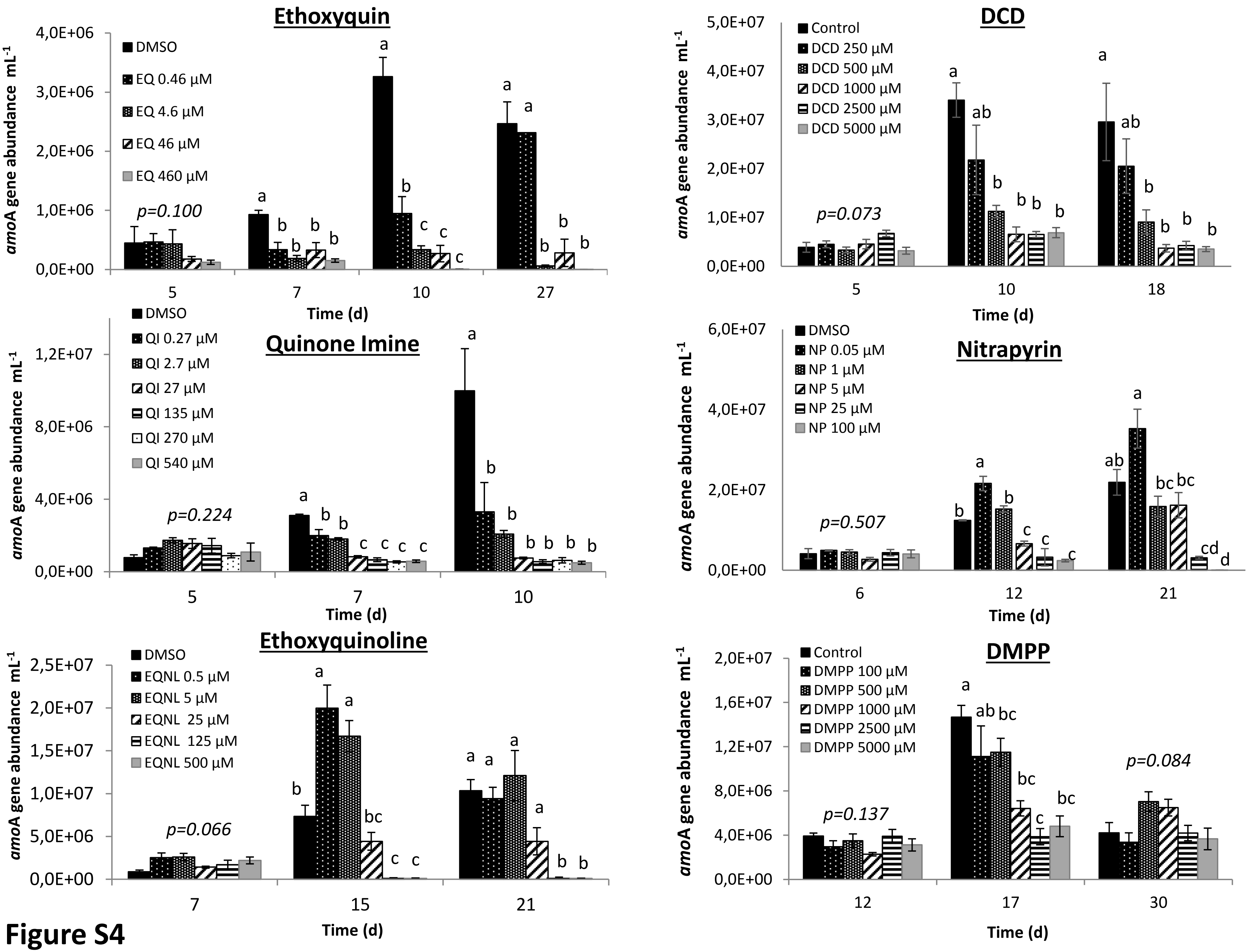
Ethoxyquin
DMSO
EQ 0.46 µM
EQ 4.6 µM
EQ 46 µM
EQ 460 µM
p=0.100
amoA gene abundance mL-1
Time (d)
DCD
Control
DCD 250 µM
DCD 500 µM
DCD 1000 µM
DCD 2500 µM
DCD 5000 µM
p=0.073
Quinone Imine
DMSO
QI 0.27 µM
QI 2.7 µM
QI 27 µM
QI 135 µM
QI 270 µM
QI 540 µM
p=0.224
Nitrapyrin
DMSO
NP 0.05 µM
NP 1 µM
NP 5 µM
NP 25 µM
NP 100 µM
p=0.507
Ethoxyquinoline
DMSO
EQNL 0.5 µM
EQNL 5 µM
EQNL 25 µM
EQNL 125 µM
EQNL 500 µM
p=0.066
DMPP
Control
DMPP 100 µM
DMPP 500 µM
DMPP 1000 µM
DMPP 2500 µM
DMPP 5000 µM
p=0.137
p=0.084

**Figure S4**

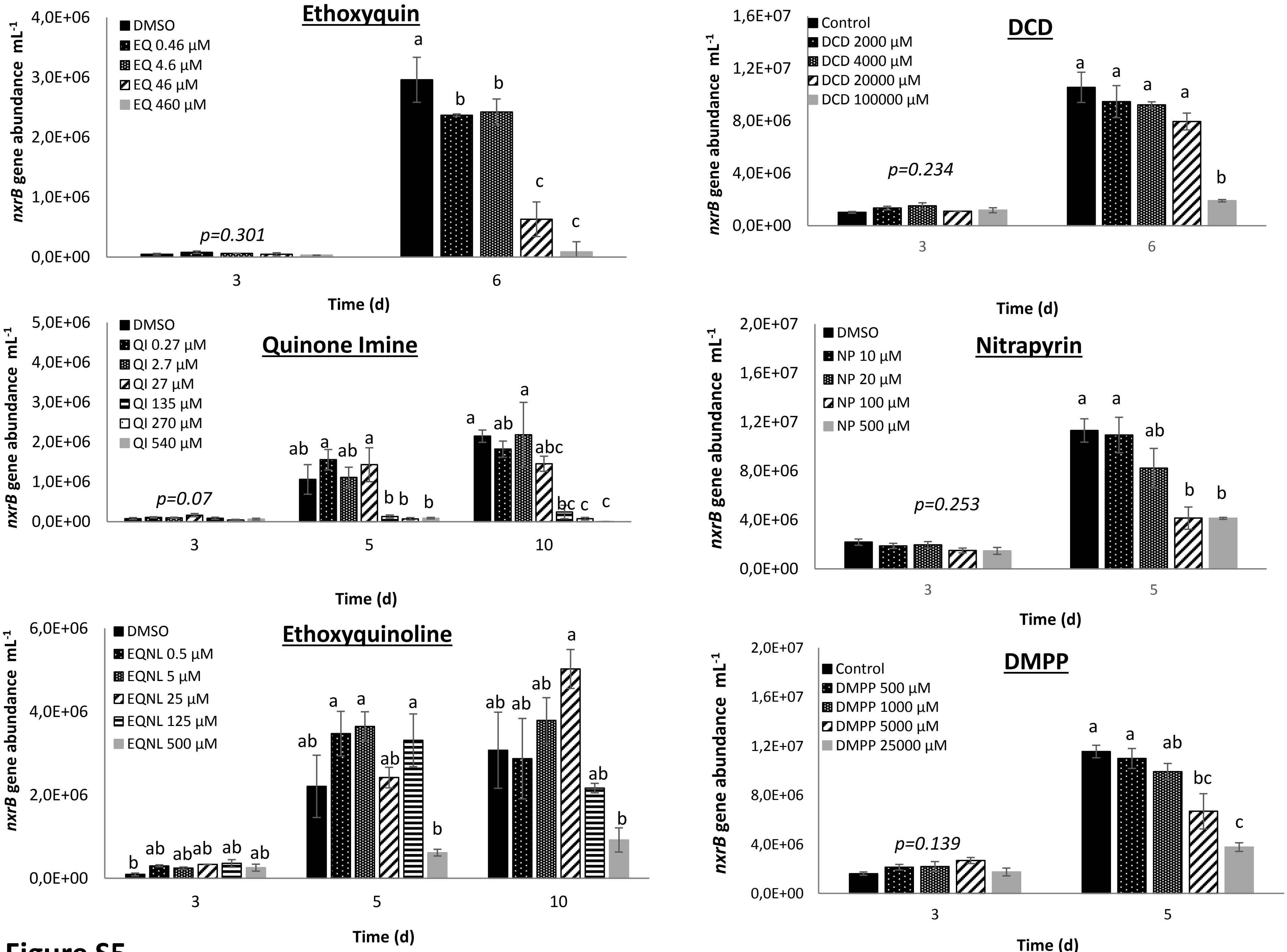
Ethoxyquin
DMSO
EQ 0.46 µM
EQ 4.6 µM
EQ 46 µM
EQ 460 µM
p=0.301
nxrB gene abundance mL-1
Time (d)
DCD
Control
DCD 2000 µM
DCD 4000 µM
DCD 20000 µM
DCD 100000 µM
p=0.234
nxrB gene abundance mL-1
Time (d)
Quinone Imine
DMSO
QI 0.27 µM
QI 2.7 µM
QI 27 µM
QI 135 µM
QI 270 µM
QI 540 µM
p=0.07
nxrB gene abundance mL-1
Time (d)
Nitrapyrin
DMSO
NP 10 µM
NP 20 µM
NP 100 µM
NP 500 µM
p=0.253
nxrB gene abundance mL-1
Time (d)
Ethoxyquinoline
DMSO
EQNL 0.5 µM
EQNL 5 µM
EQNL 25 µM
EQNL 125 µM
EQNL 500 µM
nxrB gene abundance mL-1
Time (d)
DMPP
Control
DMPP 500 µM
DMPP 1000 µM
DMPP 5000 µM
DMPP 25000 µM
p=0.139
nxrB gene abundance mL-1
Time (d)

**Figure S5**

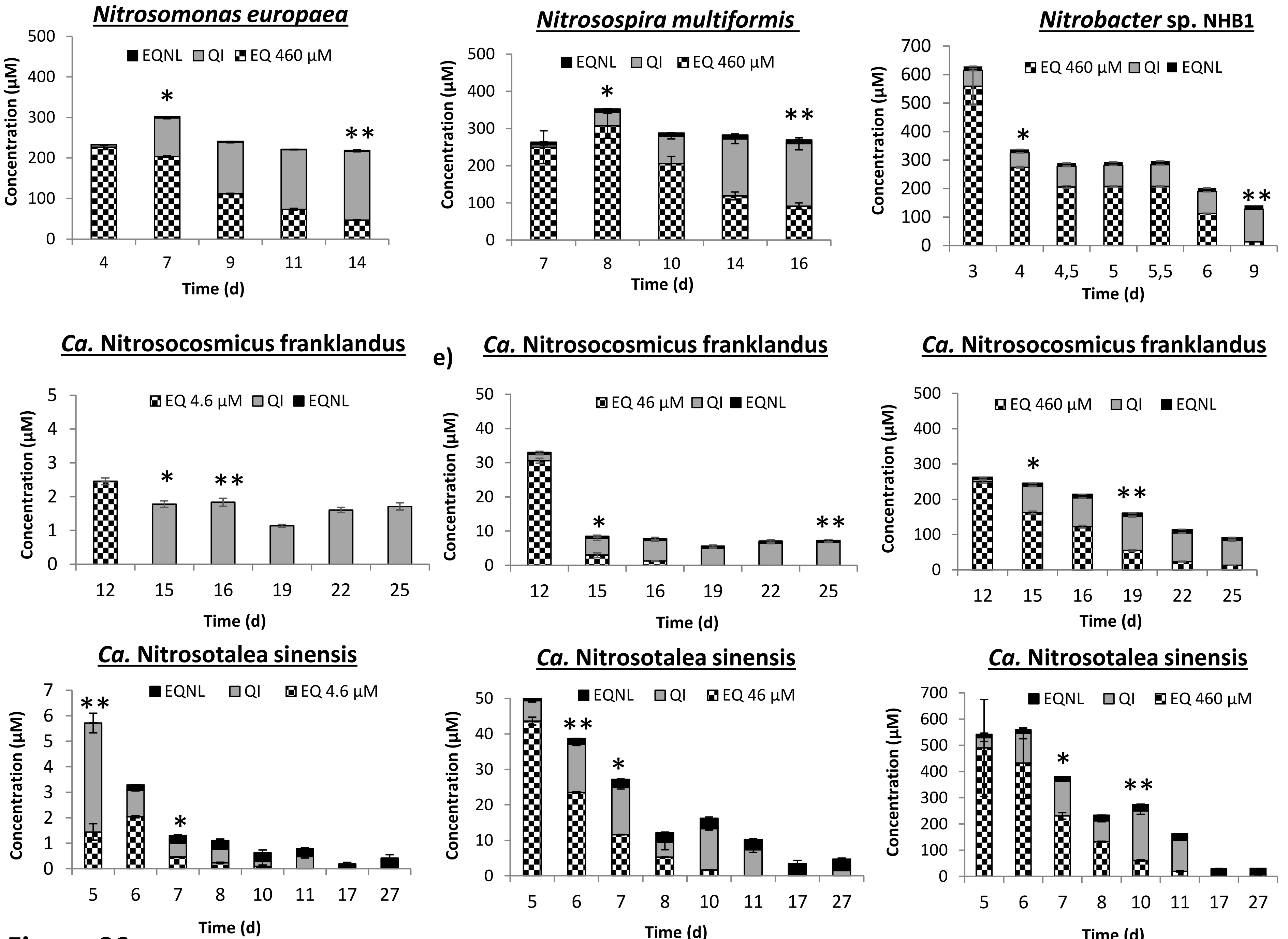
Nitrosomonas europaea
EQNL QI EQ 460 µM
Concentration (µM)
Time (d)
Nitrosospira multiformis
EQNL QI EQ 460 µM
Concentration (µM)
Time (d)
Nitrobacter sp. NHB1
EQ 460 µM QI EQNL
Concentration (µM)
Time (d)
Ca. Nitrosocosmicus franklandus
EQ 4.6 µM QI EQNL
Concentration (µM)
Time (d)
e)
Ca. Nitrosocosmicus franklandus
EQ 46 µM QI EQNL
Concentration (µM)
Time (d)
Ca. Nitrosocosmicus franklandus
EQ 460 µM QI EQNL
Concentration (µM)
Time (d)
Ca. Nitrosotalea sinensis
EQNL QI EQ 4.6 µM
Concentration (µM)
Time (d)
Ca. Nitrosotalea sinensis
EQNL QI EQ 46 µM
Concentration (µM)
Time (d)
Ca. Nitrosotalea sinensis
EQNL QI EQ 460 µM
Concentration (µM)
Time (d)

**Figure S6**

**Ethoxyquin**

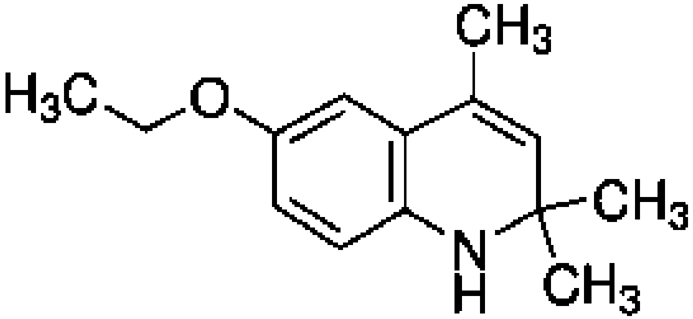


**Quinone Imine**

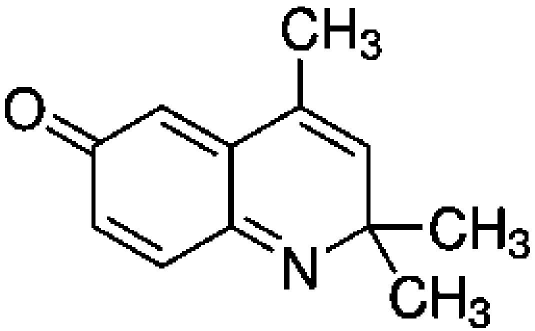


**QI**

**Ethoxyquinoline**

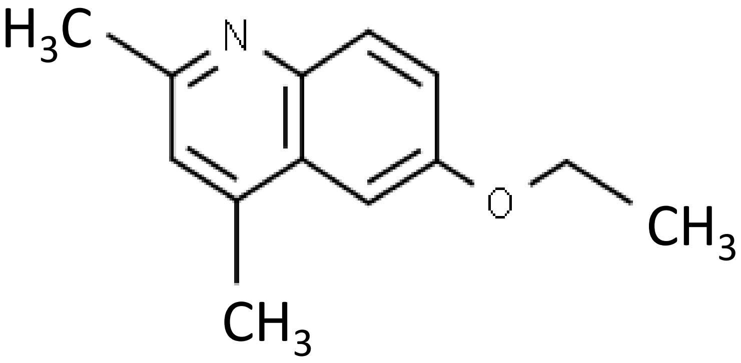


**DCD**

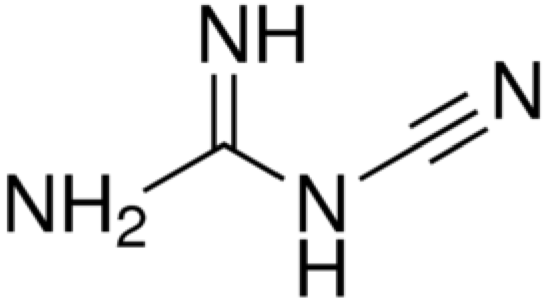


**Nitrapyrin**

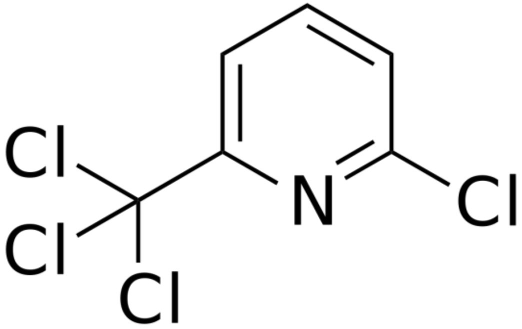


**DMPP**

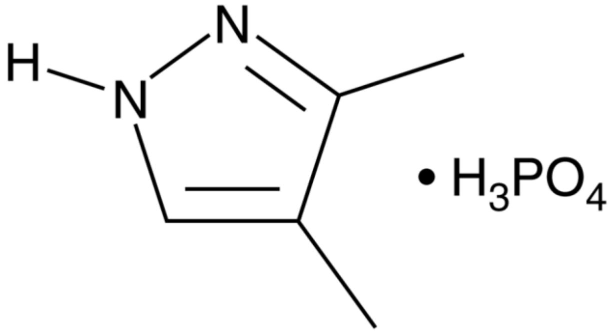

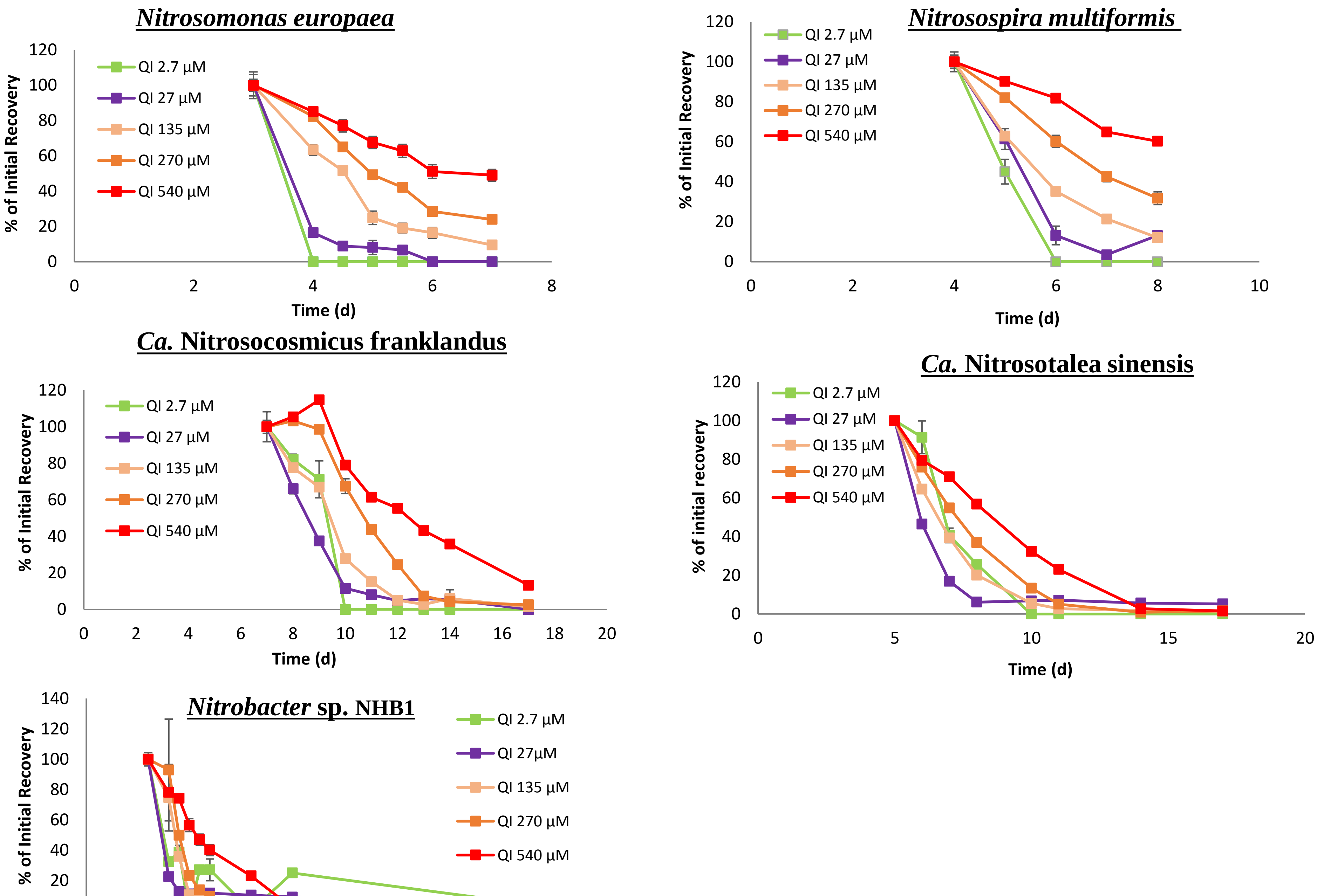
Nitrosomonas europaea
QI 2.7 µM
QI 27 µM
QI 135 µM
QI 270 µM
QI 540 µM
% of Initial Recovery
Time (d)
Nitrosospira multiformis
QI 2.7 µM
QI 27 µM
QI 135 µM
QI 270 µM
QI 540 µM
% of Initial Recovery
Time (d)
Ca. Nitrosocosmicus franklandus
QI 2.7 µM
QI 27 µM
QI 135 µM
QI 270 µM
QI 540 µM
% of Initial Recovery
Time (d)
Ca. Nitrosotalea sinensis
QI 2.7 µM
QI 27 µM
QI 135 µM
QI 270 µM
QI 540 µM
% of initial recovery
Time (d)
Nitrobacter sp. NHB1
QI 2.7 µM
QI 27µM
QI 135 µM
QI 270 µM
QI 540 µM
% of Initial Recovery
Time (d)

**Figure S8**

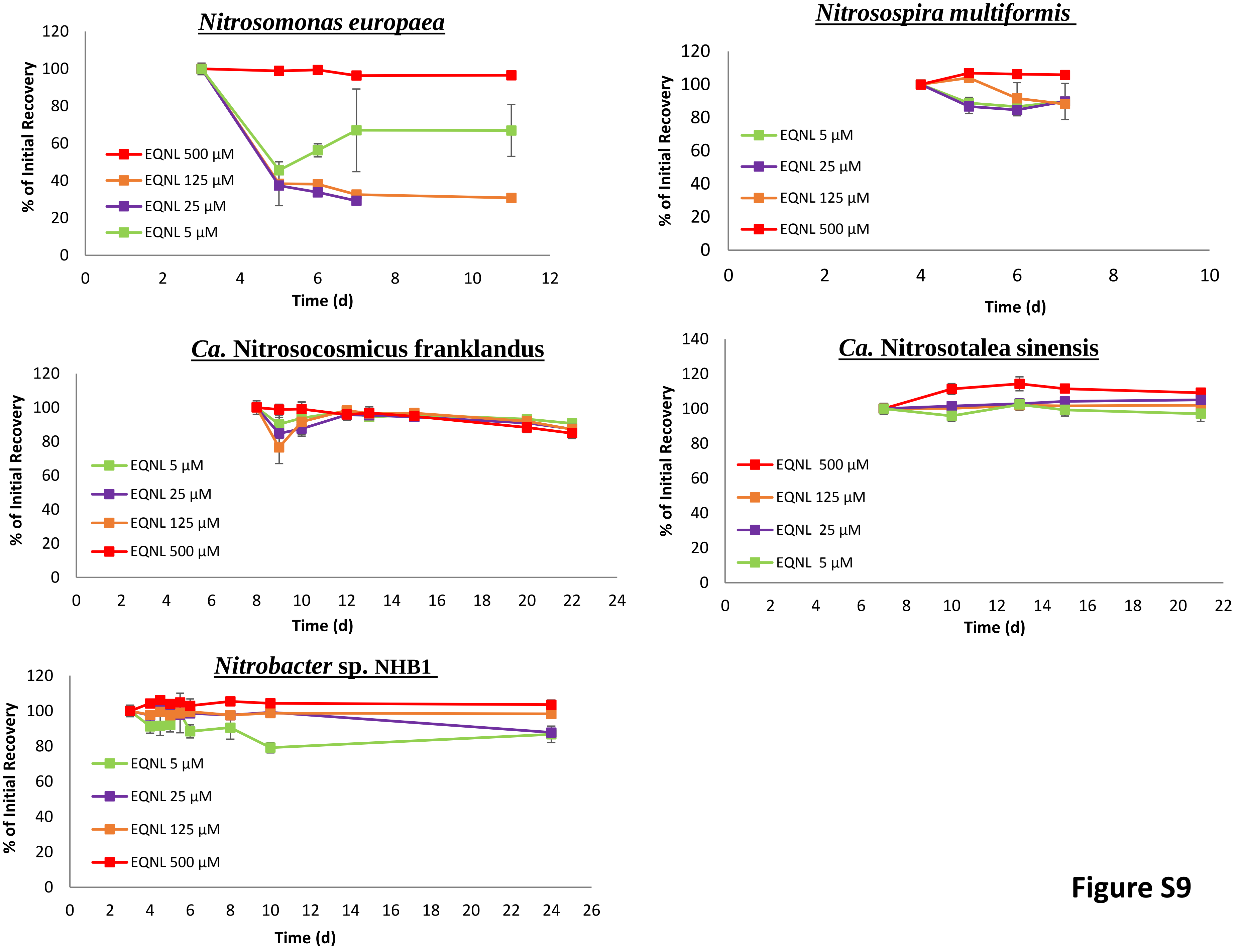
Nitrosomonas europaea
% of Initial Recovery
Time (d)
EQNL 500 µM
EQNL 125 µM
EQNL 25 µM
EQNL 5 µM
Nitrosospira multiformis
% of Initial Recovery
Time (d)
EQNL 5 µM
EQNL 25 µM
EQNL 125 µM
EQNL 500 µM
Ca. Nitrosocosmicus franklandus
% of Initial Recovery
Time (d)
EQNL 5 µM
EQNL 25 µM
EQNL 125 µM
EQNL 500 µM
Ca. Nitrosotalea sinensis
% of Initial Recovery
Time (d)
EQNL 500 µM
EQNL 125 µM
EQNL 25 µM
EQNL 5 µM
Nitrobacter sp. NHB1
% of Initial Recovery
Time (d)
EQNL 5 µM
EQNL 25 µM
EQNL 125 µM
EQNL 500 µM

**Figure S9**

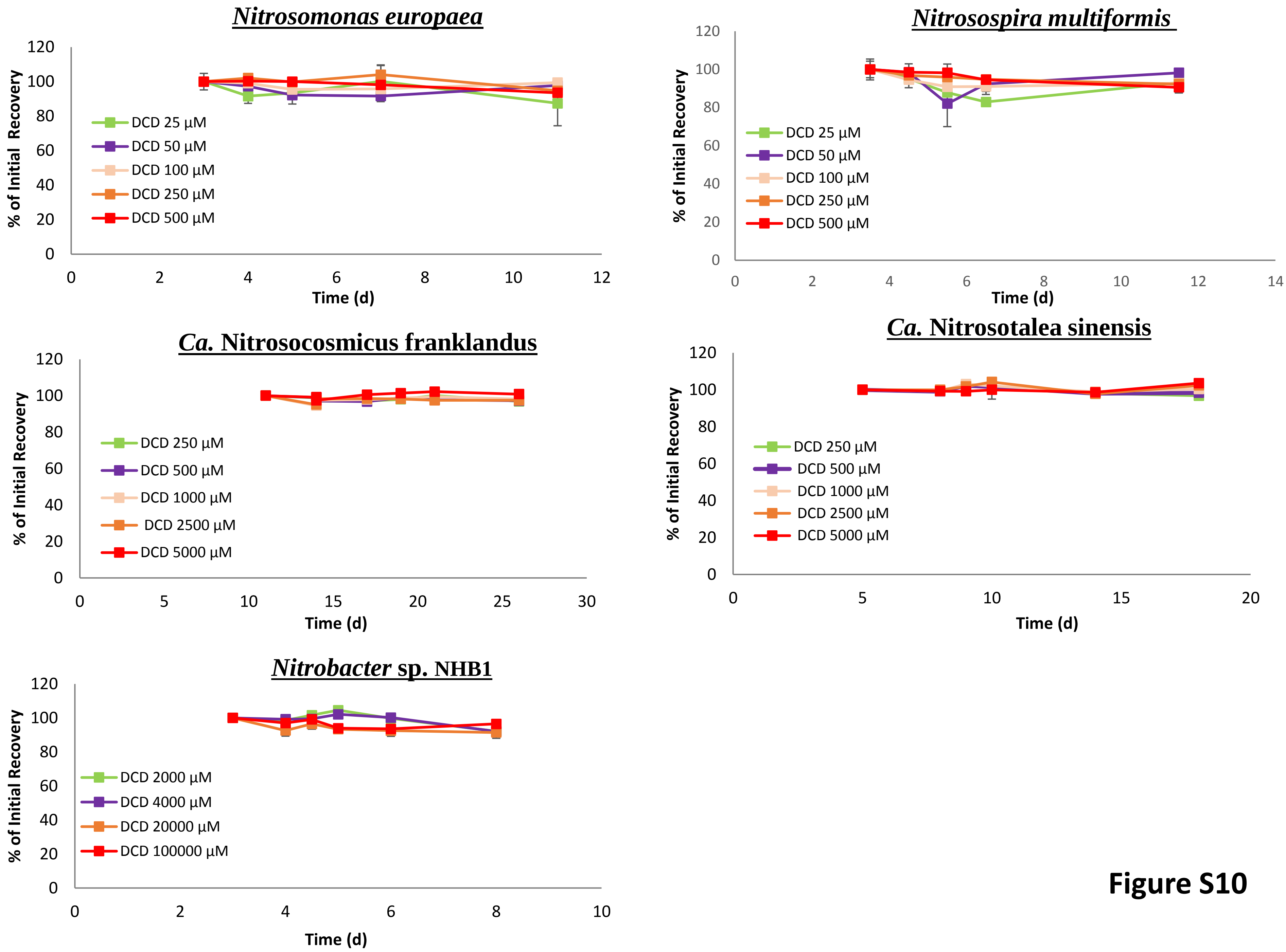

Nitrosomonas europaea
% of Initial Recovery
Time (d)
DCD 25 µM
DCD 50 µM
DCD 100 µM
DCD 250 µM
DCD 500 µM
Nitrosospira multiformis
% of Initial Recovery
Time (d)
DCD 25 µM
DCD 50 µM
DCD 100 µM
DCD 250 µM
DCD 500 µM
Ca. Nitrosocosmicus franklandus
% of Initial Recovery
Time (d)
DCD 250 µM
DCD 500 µM
DCD 1000 µM
DCD 2500 µM
DCD 5000 µM
Ca. Nitrosotalea sinensis
% of Initial Recovery
Time (d)
DCD 250 µM
DCD 500 µM
DCD 1000 µM
DCD 2500 µM
DCD 5000 µM
Nitrobacter sp. NHB1
% of Initial Recovery
Time (d)
DCD 2000 µM
DCD 4000 µM
DCD 20000 µM
DCD 100000 µM


**Figure S10**

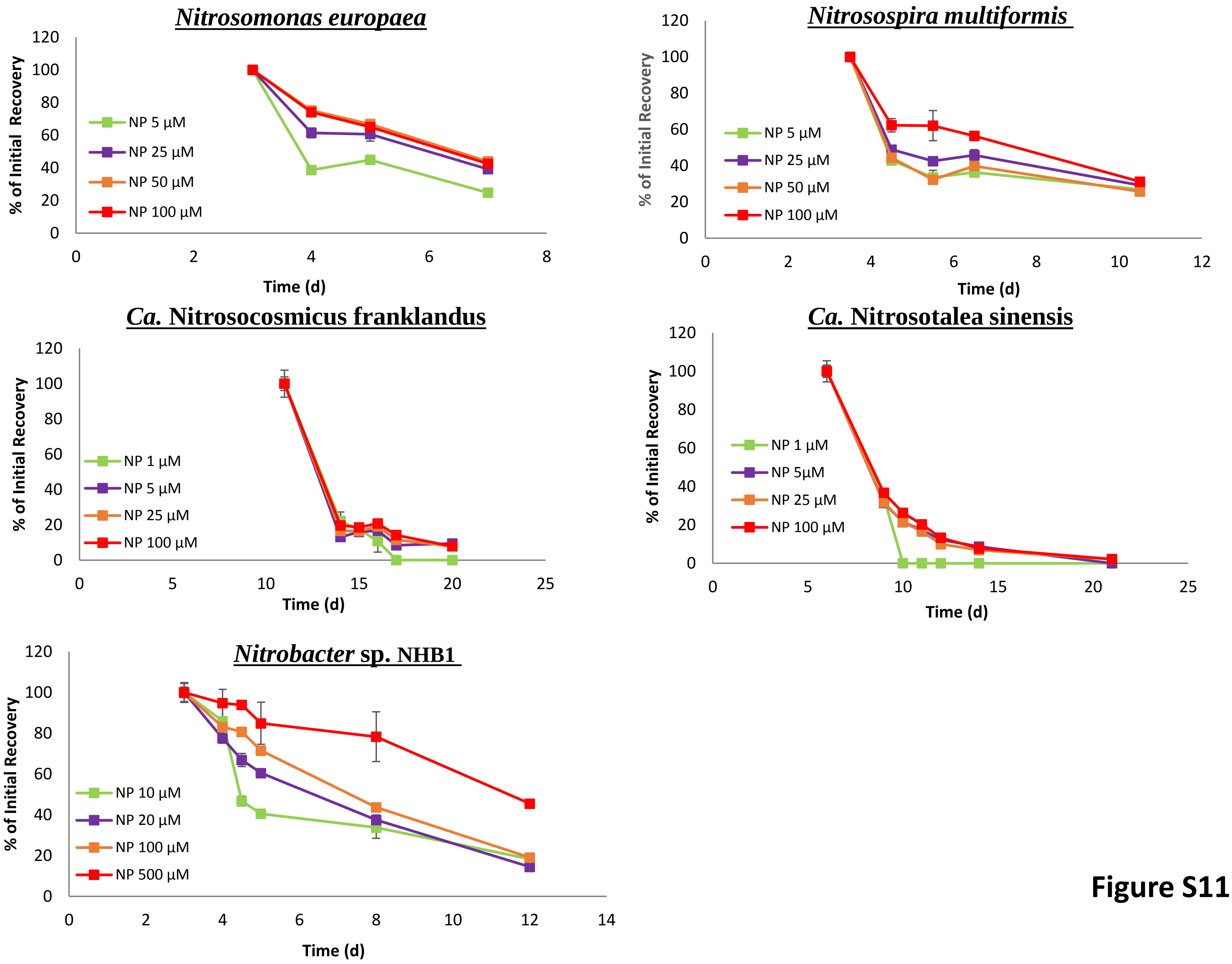
Nitrosomonas europaea
Nitrosospira multiformis
Ca. Nitrosocosmicus franklandus
Ca. Nitrosotalea sinensis
Nitrobacter sp. NHB1
% of Initial Recovery
Time (d)
NP 1 µM
NP 5 µM
NP 5µM
NP 10 µM
NP 20 µM
NP 25 µM
NP 50 µM
NP 100 µM
NP 500 µM

Figure S11

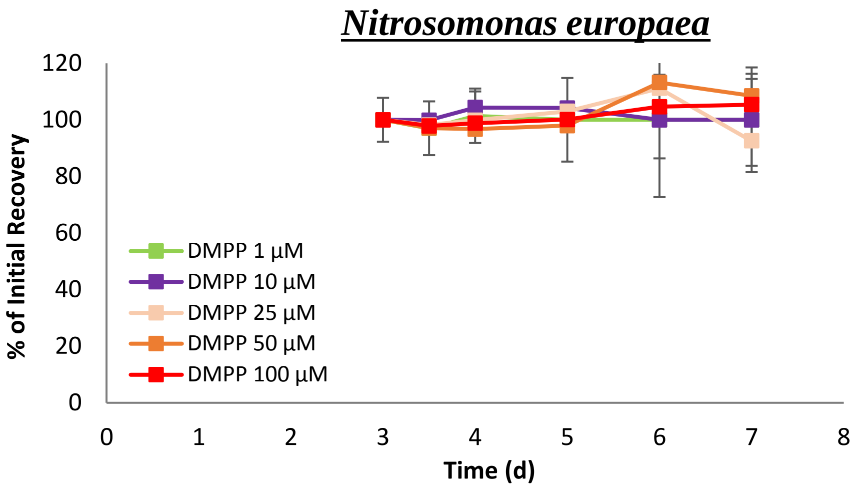

Nitrosomonas europaea
% of Initial Recovery
0
20
40
60
80
100
120
Time (d)
0 1 2 3 4 5 6 7 8
DMPP 1 µM
DMPP 10 µM
DMPP 25 µM
DMPP 50 µM
DMPP 100 µM


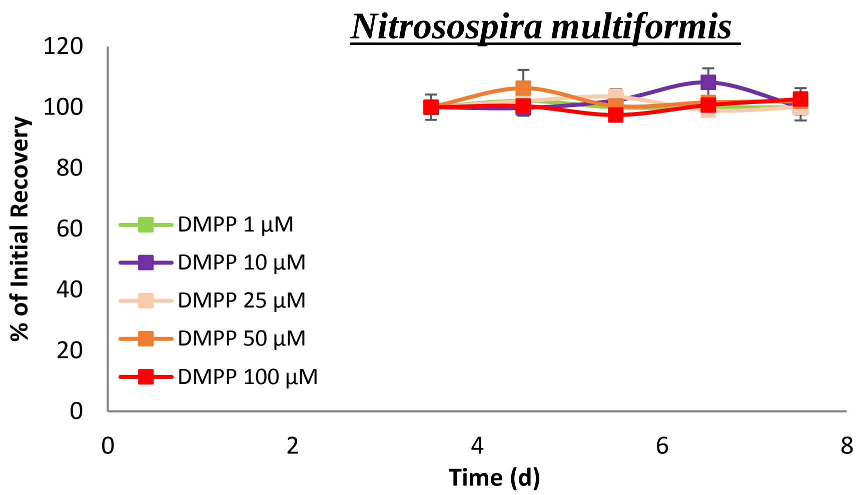

Nitrosospira multiformis
% of Initial Recovery
0
20
40
60
80
100
120
Time (d)
0 2 4 6 8
DMPP 1 µM
DMPP 10 µM
DMPP 25 µM
DMPP 50 µM
DMPP 100 µM


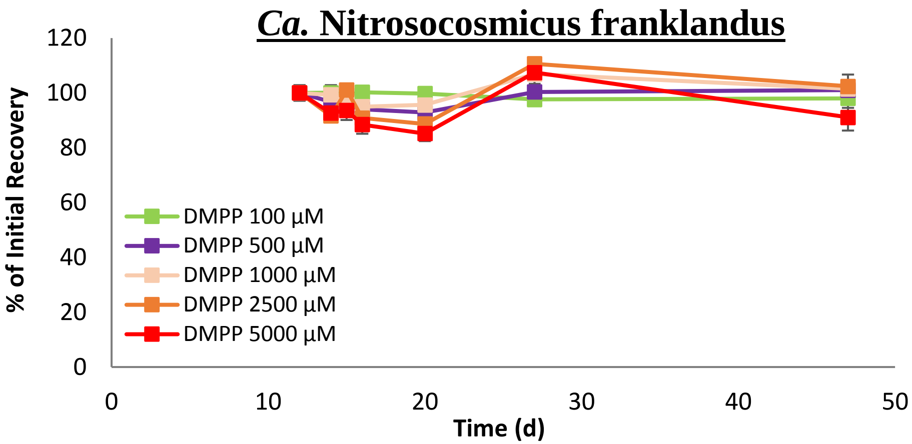

Ca. Nitrosocosmicus franklandus
% of Initial Recovery
0
20
40
60
80
100
120
Time (d)
0 10 20 30 40 50
DMPP 100 µM
DMPP 500 µM
DMPP 1000 µM
DMPP 2500 µM
DMPP 5000 µM


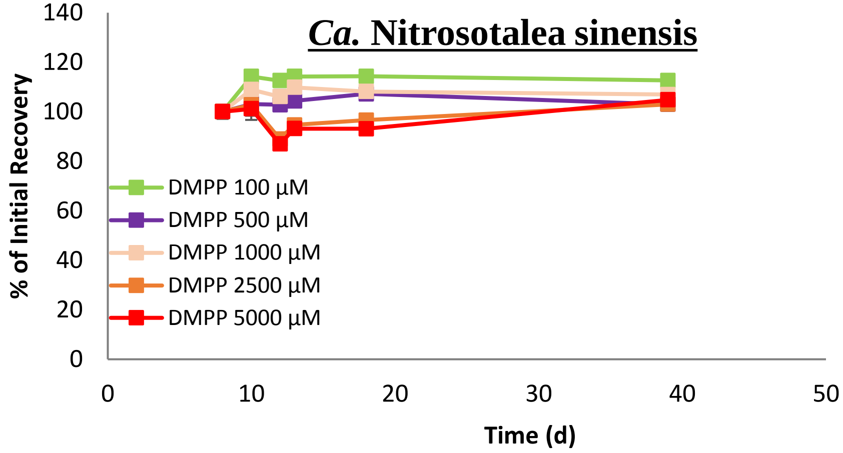

Ca. Nitrosotalea sinensis
% of Initial Recovery
0
20
40
60
80
100
120
140
Time (d)
0 10 20 30 40 50
DMPP 100 µM
DMPP 500 µM
DMPP 1000 µM
DMPP 2500 µM
DMPP 5000 µM


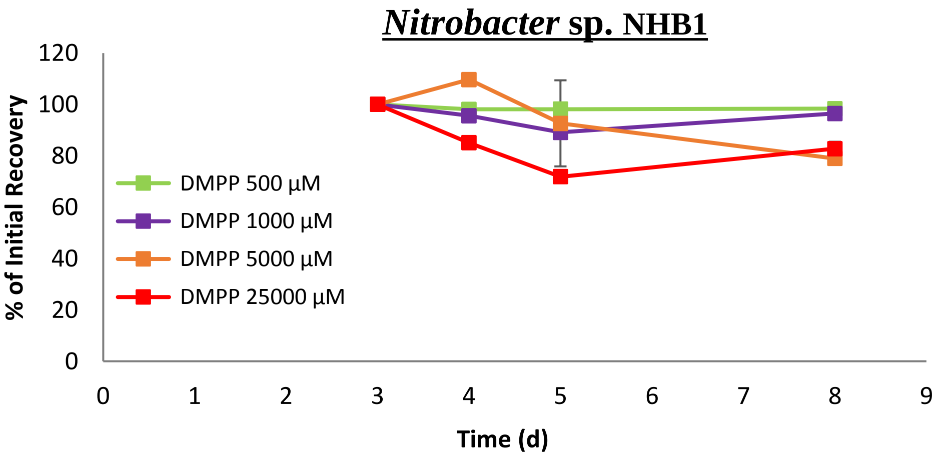

Nitrobacter sp. NHB1
% of Initial Recovery
0
20
40
60
80
100
120
Time (d)
0 1 2 3 4 5 6 7 8 9
DMPP 500 µM
DMPP 1000 µM
DMPP 5000 µM
DMPP 25000 µM


**Figure S12**
